# A transgenic line that reports CSF1R protein expression provides a definitive marker for the mouse mononuclear phagocyte system

**DOI:** 10.1101/2020.07.09.196402

**Authors:** Kathleen Grabert, Anuj Sehgal, Katharine M. Irvine, Evi Wollscheid-Lengeling, Derya D. Ozdemir, Jennifer Stables, Garry A. Luke, Martin D. Ryan, Antony Adamson, Neil E. Humphreys, Cheyenne J. Sandrock, Rocio Rojo, Veera A. Verkasalo, Werner Mueller, Peter Hohenstein, Allison R. Pettit, Clare Pridans, David A. Hume

## Abstract

The proliferation, differentiation and survival of cells of the mononuclear phagocyte system (MPS, progenitors, monocytes, macrophages and classical dendritic cells) is controlled by signals from the macrophage colony-stimulating factor receptor (CSF1R). Cells of the MPS lineage have been identified using numerous surface markers and transgenic reporters but none is both universal and lineage-restricted. Here we report the development and characterization of a novel CSF1R reporter mouse. A Fusion Red (FRed) cassette was inserted in-frame with the C-terminus of CSF1R, separated by a T2A-cleavable linker. The insertion had no effect of CSF1R expression or function. CSF1R-FRed was expressed in monocytes and macrophages and absent from granulocytes and lymphocytes. In bone marrow, CSF1R-FRed was absent in lineage-negative hematopoietic stem cells (HSC), arguing against a direct role for CSF1R in myeloid lineage commitment. It was highly-expressed in marrow monocytes and common myeloid progenitors (CMP) but significantly lower in granulocyte-macrophage progenitors (GMP). In sections of bone marrow, CSF1R-FRed was also detected in osteoclasts, CD169_+_ resident macrophages and, consistent with previous mRNA analysis, in megakaryocytes. In lymphoid tissues, CSF1R-FRed highlighted diverse MPS populations including classical dendritic cells. Whole mount imaging of non-lymphoid tissues in mice with combined CSF1R-FRed/*Csf1r*-EGFP confirmed the restriction of CSF1R expression to MPS cells. The two markers highlight the remarkable abundance and regular distribution of tissue MPS cells including novel macrophage populations within tendon and skeletal muscle and underlying the mesothelial/serosal/capsular surfaces of every major organ. The CSF1R-FRed mouse provides a novel reporter with exquisite specificity for cells of the MPS.

## Introduction

The proliferation, differentiation and survival of most tissue macrophages depends upon signals from the macrophage colony-stimulating factor receptor (CSF1R). Homozygous recessive mutations in the receptor in mice, rats and humans lead to loss of tissue macrophages and osteoclasts and pleiotropic developmental abnormalities in many organs systems (reviewed in (1)). *Csf1r* mRNa is expressed by all resident tissue macrophages (2). The transcriptional regulation of the *Csf1r* gene has been studied extensively as a model of macrophage differentiation (3). Deletion of a conserved enhancer in the first intron (Fms intronic regulatory element, FIRE) leads to selective loss of *Csf1r* expression in tissue macrophage populations including microglia in the brain (4). Transgenic reporters containing FIRE have been generated in mice (5, 6) rats (7), chickens (8, 9) and sheep (10) and in each species highlight the location and abundance of macrophage populations in every organ. However, there are caveats to the utility and interpretation of these *Csf1r* reporters. Firstly, based upon the phenotype of the *Csf1r*_ΔFIRE/ΔFIRE_ mice, we identified regulatory elements outside of the transgene construct, so that the absence of reporter gene detection in particular cells may be difficult to interpret (4). In the chick, we found that Kupffer cells expressed *Csf1r* mRNA and protein but did not express the *Csf1r*-mApple reporter transgene (11). Conversely, mouse and human granulocytes express *Csf1r* mRNA but do not translate the protein (12) and the *Csf1r* promoter also has detectable activity in B lymphocytes, which share expression of the transcriptional regulator PU.1 (13, 14). Accordingly, the *Csf1r* reporter transgenes in mouse, rat and chicken were detectable in both granulocytes and B cells. In many tissues this is not a major issue; the macrophages have a characteristic location and stellate morphology (15, 16). But in lymphoid tissues, and mucosal surfaces and inflammatory sites, it would be desirable to separate these cell types.

There have been a number of reports on expression of CSF1R in non-hematopoietic cells including neurons and neuronal progenitors in the brain and epithelial cells in the gut and kidney (17–19). In each case, the conclusion depends upon immunolocalization with poorly-characterized polyclonal antibodies or conditional reporter transgenes and was not supported by subsequent analyses (1). In the brain, the mouse *Csf1r*-EGFP reporter transgene and the protein detected by anti-CD115 antibody were expressed exclusively in microglia (20, 21). A further argument against functional expression of CSF1R in non-hematopoietic cells follows from the complete rescue of the pleiotropic impacts of *Csf1r* knockouts in mice (22) by postnatal adoptive transfer of wild-type bone marrow (BM) cells.

By contrast to surface markers such as F4/80, CD11b/c, CD163, CD68, CD169, CD64, CD206, CX3CR1, LYVE1 and MHCII, which are expressed independently by subsets of tissue MPS cells (2, 15), CSF1R is a potential universal marker for cells of the mononuclear phagocyte system (MPS). There are available monoclonal antibodies against mouse CSF1R protein (CD115). Two different antibodies have been described that block CSF1 binding to the mouse receptor and can deplete tissue macrophages (23, 24). Anti-CD115 antibodies can detect cell surface expression on at least some myeloid progenitors in the BM (25) and on isolated blood monocytes (6) and can compete for binding of labelled CSF1 protein to tissue macrophages *in vivo* (5). However, CSF1R protein is not readily detected on most tissue macrophages by immunohistochemistry using monoclonal anti-CD115. The lack of detection is most likely a consequence of competition with endogenous ligand (the available antibodies compete for CSF1 binding) and the down-regulation of the receptor from the cell surface by ligand. CSF1 is internalized by receptor-mediated endocytosis and both receptor and ligand are degraded (26). Accordingly, signaling requires continuous synthesis of new receptors. The surface receptor is also subject to rapid proteolytic cleavage in response to signals that activate macrophages (27, 28) and it is likely cleaved from the surface directly or indirectly during tissue disaggregation to isolate macrophages. All cell isolation procedures lead to activation of inflammatory genes in macrophages (2).

Classical dendritic cells (cDC) were originally defined by their unique ability to present antigen to naïve T cells. Class II MHC_+_ migratory monocytes, monocyte-derived cells and subsets of resident tissue macrophages also possess active antigen presentation activity (29–31). cDC and monocytes share a committed progenitor in the BM that expresses CSF1R(32). cDC have been classified separately from macrophages in different tissues and contexts based upon the lack of certain surface markers, including CD64 (Fcgr1) and MERTK (33). One subset of mouse cDC, termed cDC2, share many markers with monocytes and macrophages, including high expression of *Csf1r* mRNA and of *Csf1r* reporter transgenes (5–7). cDC retain expression of the growth factor receptor FLT3, co-expressed with CSF1R on progenitors (25) and lymphoid tissue cDC are depleted in FLT3 or FT3L knockouts. However, CSF1R signaling can rescue the impacts of *Flt3* null mutation on lymphoid tissue cDC numbers (34) and CSF1 treatment can expand their numbers in the spleen (35). By contrast, cDC2 in non-lymphoid tissues depend upon CSF1R (36, 37) and all cells in non-lymphoid tissues expressing the *Csf1r* reporter transgene were completely depleted by anti-CSF1R antibody (23). We were therefore interested in whether DC defined by current markers express CSF1R protein.

In this study, we aimed to overcome the rapid turnover of surface CSF1R and ectopic expression in promoter-based transgenics by integrating a reporter gene into the mouse *Csf1r* locus. Knock-in reporters, for example in the *Cx3Cr1*(38) and *Ccr2* loci (39) have been widely-utilized in the study of macrophage biology, but in each case the endogenous gene is knocked out. Although heterozygous mutation of *Csf1r* has no overt phenotypic impact in mouse, rat or human, there is no dosage compensation (1). Given the central role of CSF1R in macrophage homeostasis, a knock-out insertion is undesirable. One alternative is to insert a cassette including an internal ribosome re-entry site (IRES) downstream of the stop codon. This approach was used to generate a *Ms4a3* locus reporter, that tags cells derived from myeloid progenitors and apparently separated monocyte-derived cells from cDC in tissues (40). To target microglia, Ruan et al (41) inserted a tdTomato reporter with a cleavable peptide linker between the coding sequence and 3’UTR of the *Tmem119* locus. Masuda *et al.* (42) inserted the same reporter into the *Hexb* locus. Here we took a similar approach and inserted a Fusion-Red (FRed) cassette with cleavable linker in-frame into the 3’ end of the mouse *Csf1r* locus. Analysis of this novel line confirms that CSF1R protein expression is restricted to macrophages and their progenitors and provides a universal functional marker for cells of the MPS lineage. Imaging of this reporter gene provides a unique picture of the extent of the MPS in tissues.

## Results

### Development and validation of a strategy to generate a translational CSF1R reporter

To overcome the limitations of the current *Csf1r* reporter transgenes we aimed to generate a transgene that would accurately mirror protein expression. The ideal outcome would place a reporter cassette in-frame with the CSF1R protein to generate a fusion protein that would mark the plasma membrane and allow analysis of intracellular trafficking. We previously expressed functional CSF1R from several species in transfected cells with a V5-His C terminal tag (43, 44) and anticipated that it could be possible to create a CSF1R-red fluorescent protein (RFP) fusion. To enable double imaging with EGFP reporter lines we chose Fusion Red (FRed). This protein has been tested as a fusion partner in multiple applications and has the additional advantage of being resistant to photobleaching (45). A CSF1R-FRed fusion allele was generated by CRISPR-mediated recombination in embryonic stem cells (ESC) and expression was analyzed in ESC-derived macrophages (ESDM) as described previously (4). We confirmed the successful targeting of one allele in ESC, which did not prevent the generation of ESDM. However, there was no detectable FRed expression. As an alternative, we tested the insertion of a cleavable linker. For this purpose, we used the T2A peptide also used in the recent targeting of *Hexb* (42). T2A was originally identified in picornavirus (46) and enables the generation of multiple protein from a single viral mRNA. 2A oligopeptides mediate ribosomal skipping which appears as “self-cleavage” with stoichiometric expression of two or more translation products. Self-cleaving peptide linkers have been used in multicistronic vectors *in vitro* (47) and *in vivo* (48, 49). FRed was absent from transfected ESC but was detected in ESDM by flow cytometry and direct imaging (not shown). On this basis we proceeded to generate CSF1R-FRed mice as outlined in **Figure 1**.

**Figure 1.**
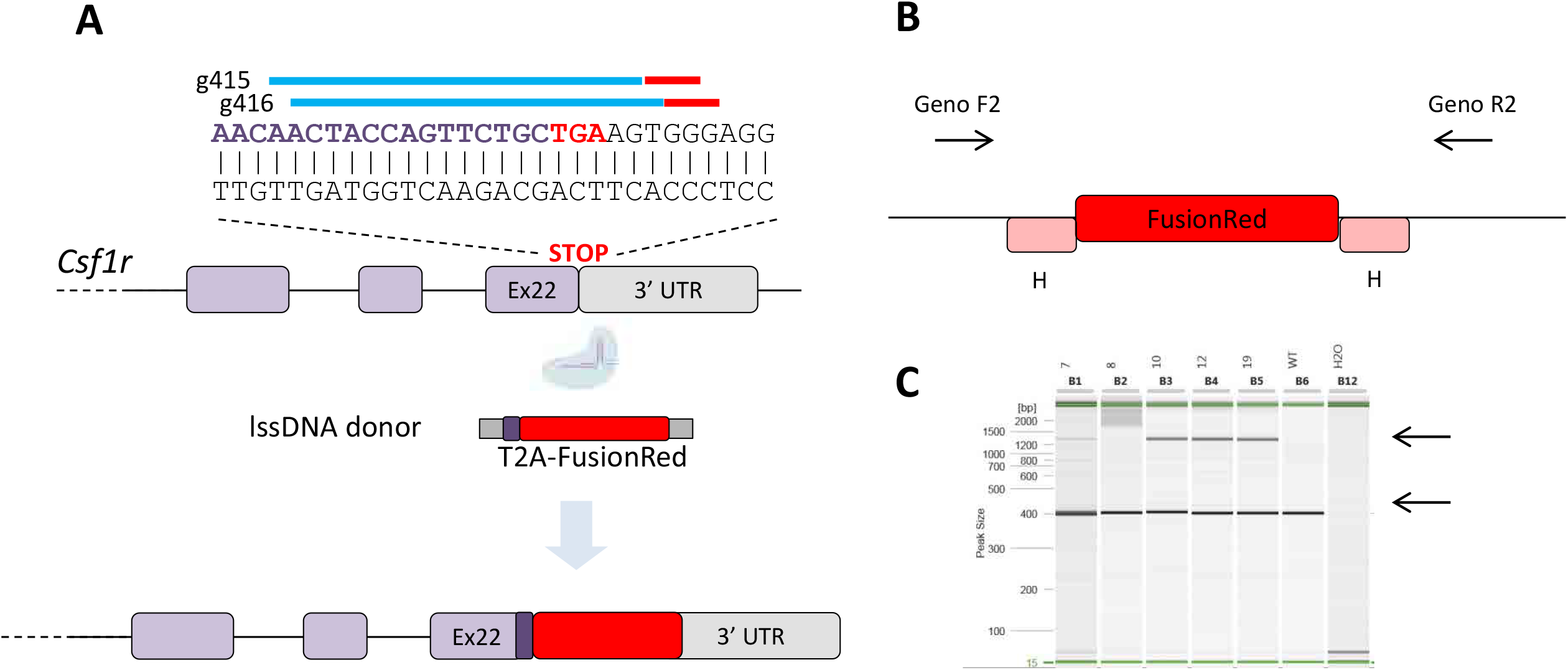
Design and generation of CSF1R-FRed transgene. Panel A shows the schematic of the insertion of T2A-Fusion Red into the mouse *Csf1r* locus with the guides and overall strategy as described in Materials and Methods. Panel B shows the location of the PCR primers and Panel C the detection of 3 independent founders with the desired insertion which was sequence verified.

### Insertion of the FRed cassette does not compromise CSF1R expression or macrophage differentiation

The *Csf1r* knockout on the C57Bl/6 background is severely compromised and few homozygotes survive to weaning (21). By contrast, the CSF1R-FRed mice were healthy and fertile, and there was no apparent effect on postnatal growth or any overt phenotype in homozygotes. We first confirmed that the FRed reporter was expressed in macrophages and their progenitors and correlated with CSF1R expression. **Figure 2** compares expression of the reporter and CSF1R (CD115) in BM cells from adult WT, heterozygous and homozygous CSF1R-FRed mice. In fresh BM, FRed expression was restricted to CD115_+_ cells **(Figure 2A).** with the exception of a minor population of hematopoietic progenitors analysed further below. In homozygotes, FRed was increased proportionate to gene dose as expected but CD115 expression was unaffected **(Figure 2B/C).** To confirm that the FRed insertion did not impact CSF1R function, we cultured BM from WT, heterozygous and homozygous CSF1R-FRed mice in CSF1 to generate BM-derived macrophages (BMDM). The yield of BMDM was unaffected by the insertion. The FRed reporter was expressed homogeneously in BMDM from heterozygous mice and around 2-fold higher in the homozygotes **(Figure 2D).** Unless otherwise stated, all analysis in this study was carried out on CSF1R-FRed heterozygotes but for some applications a homozygote might have greater utility.

**Figure 2.**
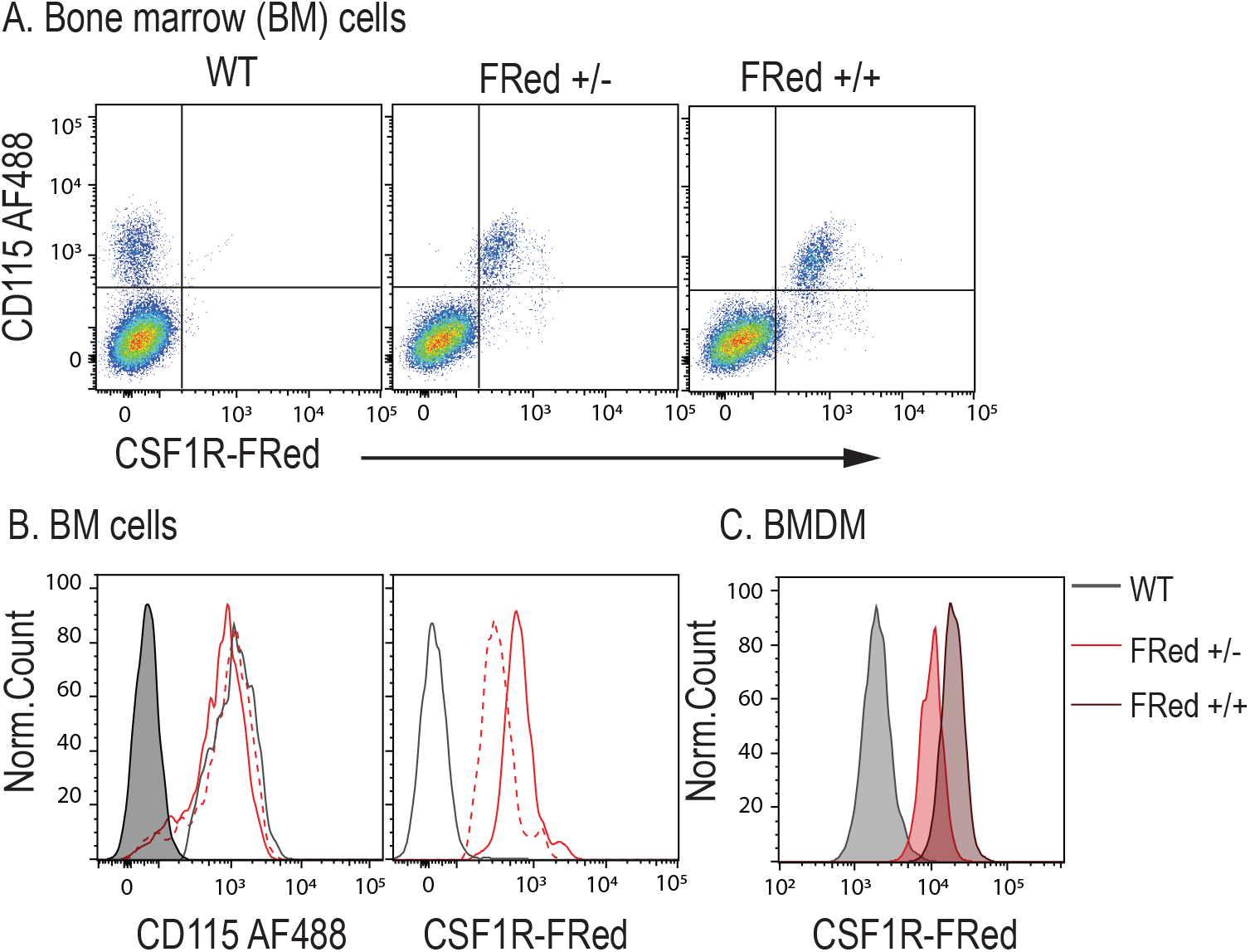
Expression of CSF1R-FRed in bone marrow (BM) and bone marrow-derived macrophages (BMDM). Panel A shows the relationship between CSF1R-FRed and surface CSF1R (CD115) on unfractionated BM cells from adult mice in WT (C57Bl/6 control), heterozygous (FRed +/−) and homozygous FRed (+/+) mice. Panel C shows histograms of the same profiles, with fill profile WT, dotted line (FRed +/−) and filled line FRed (+/+). Note that the expression of the two markers is correlated and there is a right shift in FRed in the homozygote. In Panel C, BMDM were grown from WT, FRed +/− and FRed +/+ BM cells as described in Materials and methods. The cells were harvested after 7 days.

Previous studies reported that *Csf1r* mRNA is first detectable is early mouse myeloid precursors on which surface CSF1R was not detectable (50). CSF1R is expressed on the surface of committed macrophage-dendritic (MDP) and monocyte progenitors (25). The question of CSF1R expression by progenitors is key to the issue of whether CSF1 is instructive or selective in lineage commitment. It has been suggested that CSF1 instructs lineage fate in HSC by inducing the key transcriptional regulator PU.1 (51, 52). This proposal is difficult to reconcile with the fact that *Csf1r* promoter activity is stringently dependent upon PU.1 (14) and with analysis of progenitors by scRNA-seq (53), which associated *Csf1r* mRNA with committed progenitors. CSF1R-FRed provides a unique marker to address this question. Aside from stem cell and committed progenitors, BM contains monocytes and multiple distinct resident macrophage populations (54). The results of flow cytometry on isolated BM cells are summarized in **Figure 3.** Consistent with *Csf1r* mRNA expression data (50, 53), CSF1R-FRed expression was not detected in HSC or MPP and present only within a small subset of Lin-CD48_+_ and Lin-cKit_+_Sca1-progenitors **(Figure 3A)**. Committed progenitors (CLP, CMP, GMP and MEP) were separated based upon surface markers according to the hierarchical model of Akashi *et al.* (55). No FRed expression was detected in CLP **(Figure 3B)**. Surprisingly, although the two populations were uniformly-positive, the CSF1R-FRed signal was considerably higher in CMP than in GMP. Signal was also detected in MEP.

**Figure 3.**
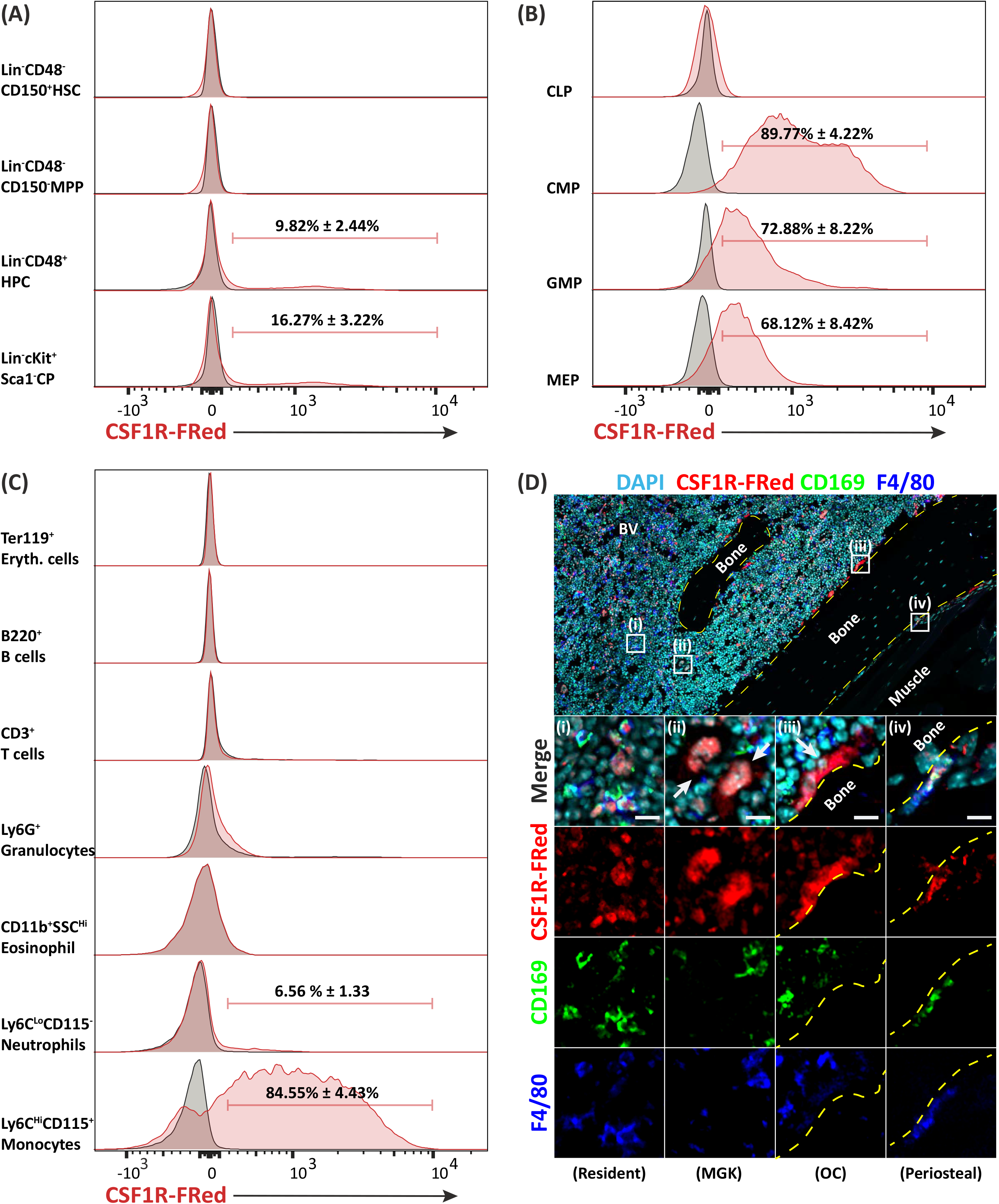
Expression of CSF1R-FRed in bone marrow. Femoral bone marrow cells were harvested from adult wildtype and CSF1R-FRed +/− mice and analysed by flow cytometry using the markers indicated and gating strategies shown in **Figure S5** and **Figure S6.** Panel A shows the defined stem cell populations, Panel B the committed progenitors and Panel C the mature leukocytes. In each case, the grey profile is the non-transgenic control and the red profile is the CSF1R-FRed mouse. The results are representative of three mice of each genotype. Panel D shows the localization of CSF1R-FRed in femoral bone marrow. Bone was fixed and decalcified as described in Material and Methods and stained for each of the markers shown. CSFR-FRed was detected using anti-RFP antibody. The staining highlights expression of CSF1R-FRed in F4/80_+_/CD169_+_ resident macrophages within hematopoietic islands and on the periosteal surface, and in megakaryocytes (MGK) and F4/80_−_/CD160_−_ osteoclasts (OC).

In flow cytometry analysis of lineage_+_ cells in BM suspensions, CSF1R-FRed was restricted to Ly6_Chi_, CD115_+_ monocytes **(Figure 3C).** By contrast to *Csf1r* reporter transgenes, granulocytes, that express *Csf1r* mRNA at high levels but do not express surface CD115 or CSF1R protein detectable by Western blot (12), were CSF1R-FRedneg. Similarly, B cells did not express CSF1R-FRed, indicating that CSF1R protein regulation is controlled by elements outside the core *Csf1r* promoter and enhancer (5).

Resident macrophage subpopulations in BM associated with hematopoietic islands, as well as osteomacs lining bone surfaces, express Siglec1 (CD169) as well as F4/80 (56) and bone-resorbing osteoclasts are also CSF1R-dependent (7). Not all of these mature myeloid cells can be reliably isolated using bone marrow flushing and anatomical location is needed for accurate identification. To complement the flow cytometry, we stained sections of bone marrow using anti-RFP antibody, CD169 and F4/80 (**Figure 3D**). CSF1R-FRed was detected in all F4/80_+_ and CD169_+_ cells forming the centre of hematopoietic islands and lining the surface of bone (osteomacs) (57) and was also detected in multinucleated osteoclasts which are F4/80_−_. We (unpublished) and others (58) have noted that *Csf1r* reporter transgenes are also expressed by megakaryocytes and their CD41_+_ progenitors in adult mouse BM, but expression of CSF1R protein has not been reported. The sections of BM show clear expression of CSF1R-FRed in the majority of megakaryocytes *in situ*.

Peripheral blood contains two populations of blood monocytes, distinguished by the level of expression of Ly6C. This is a differentiation series dependent upon CSF1R signalling. Both populations clearly express *Csf1r* mRNA and bind anti-CD115 (23, 59), albeit at relatively low levels by comparison to resident tissue macrophages (2). The Ly6C_lo_ population had somewhat higher expression of the *Csf1r*-EGFP transgene (23) but the two populations were not distinguished by *Csf1r*-mApple (5). **Figure 4A** shows the profile of expression of CSF1R-Fred in blood leukocytes. As observed in BM, there was no detectable expression of the FRed reporter in lymphocytes or granulocytes whereas both monocyte populations were clearly positive with expression marginally higher in the Ly6C_lo_ population.

**Figure 4.**
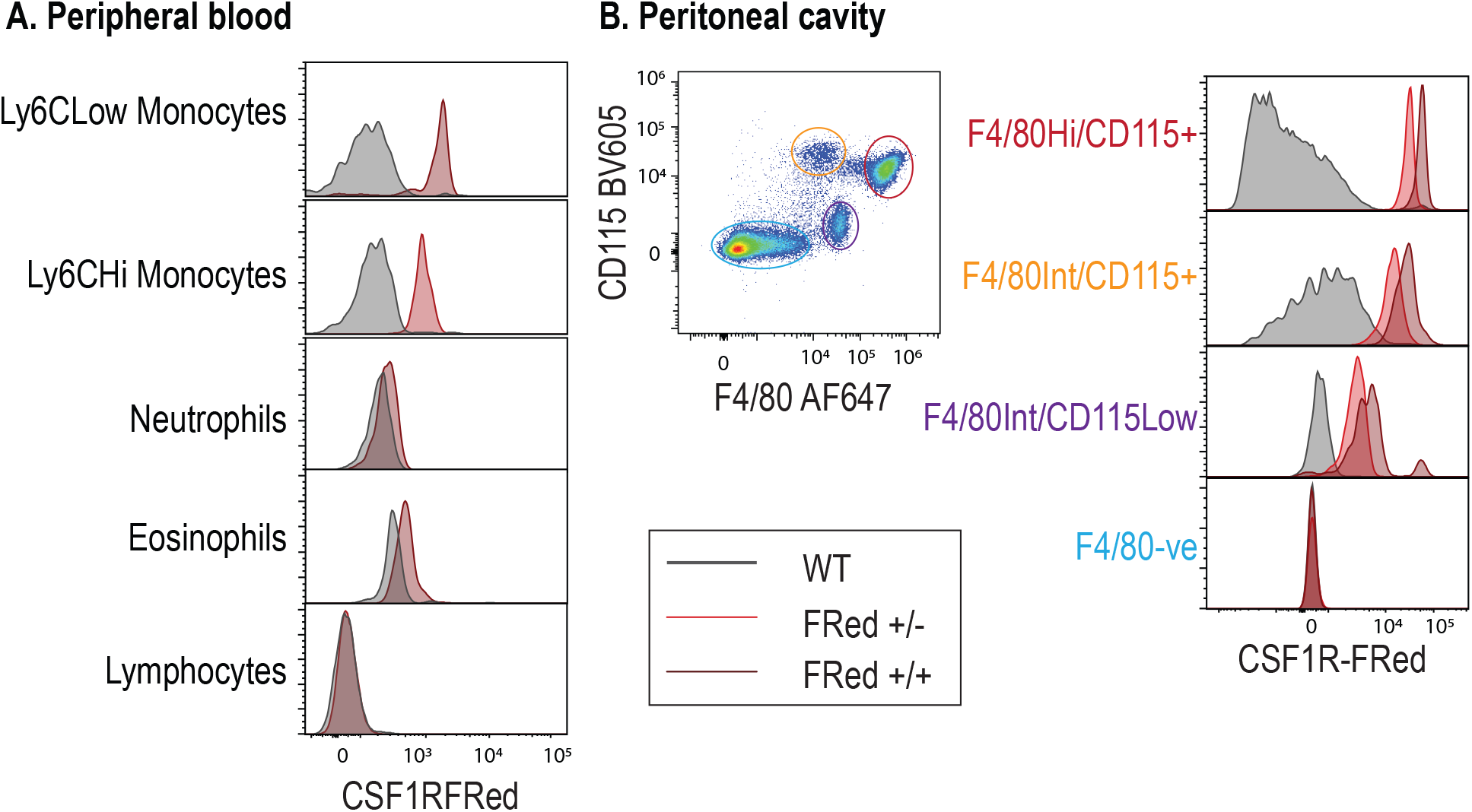
CSF1R-FRed expression in peripheral blood and peritoneal cavity leukocytes. Panel A. Peripheral blood leukocytes were isolated from WT and CSF1R-FRed +/− mice and analysed by Flow Cytometry using gating strategies shown in **Figure S7.** Red histogram show expression in the CSF1R-FRed mice. Panel B. Peritoneal lavage cells were isolated from WT, FRed +/- and FRed +/+ and separated into 4 populations based upon F4/80 and CD115 as shown. Note the increased expression in FRed +/+ mice.

In the peritoneal cavity, the major F4/80_Hi_/CD115_+_ and minor F4/80_Int_/CD115_+_ macrophage populations expressed similar levels of FRed; whereas low FRed expression was observed in F4/80_Int_/CD115_-ve_ cells (**Figure 4B**). The resident peritoneal macrophage population is very sensitive to anti-CSF1R treatment (23) and was selectively depleted in a *Csf1r* hypomorphic mutant (4). However, as in BM, homozygosity for the CSF1R-FRed allele did not affect peritoneal macrophage abundance or labelling with F4/80/CD115 and the reporter expression was approximately 2-fold higher in homozygotes compared to heterozygotes (**Figure 4B)**.

### Mononuclear phagocyte populations of lymphoid tissues

The spleen contains multiple resident macrophage and dendritic cells populations in the red pulp, the marginal zone (MZ), T cell areas and B cell areas (60). There is also a population of undifferentiated monocytes that can be recruited to inflammatory sites (61). Lymph nodes (LN) likewise contain multiple functional MPS subpopulations in the subcapsular sinus, medullary sinus, medullary cords, T cell areas and germinal centres (62, 63). Many of these populations lack expression of surface markers such as F4/80 and are susceptible to fragmentation during tissue disaggregation (62). **Figures 5A-F** shows co-localization of myeloid (CD169, CD68, F4/80) and lymphoid (B220, CD3) markers in section of spleen and lymph node. In the spleen, there was a complete overlap with F4/80 in the red pulp. In the MZ, CD169_+_ metallophils expressed CSF1R-Fred, as did a separate stellate population of F4/80_−_ macrophages, the outer MZ macrophages (64). Expression was excluded from B220 and CD3-positive lymphocytes. Multi-lineage flow cytometry analysis of disaggregated spleen cells confirmed the expression of CSF1R-FRed in Ly6_Chi_ monocytes and exclusion from non-MPS populations (**Figure 5D**). Isolated CD11c_hi_/F4/80_lo_ presumptive cDC were positive but heterogeneous for detectable CSF1R-FRed; consistent with detection of cells with varying levels of the reporter in the stellate interdigitating cells within the T cell areas of spleen **(Figure 5B).** The CSF1R-FRedlo cells within T cell areas were positive for CD68 **(Figure 5B).**

**Figure 5.**
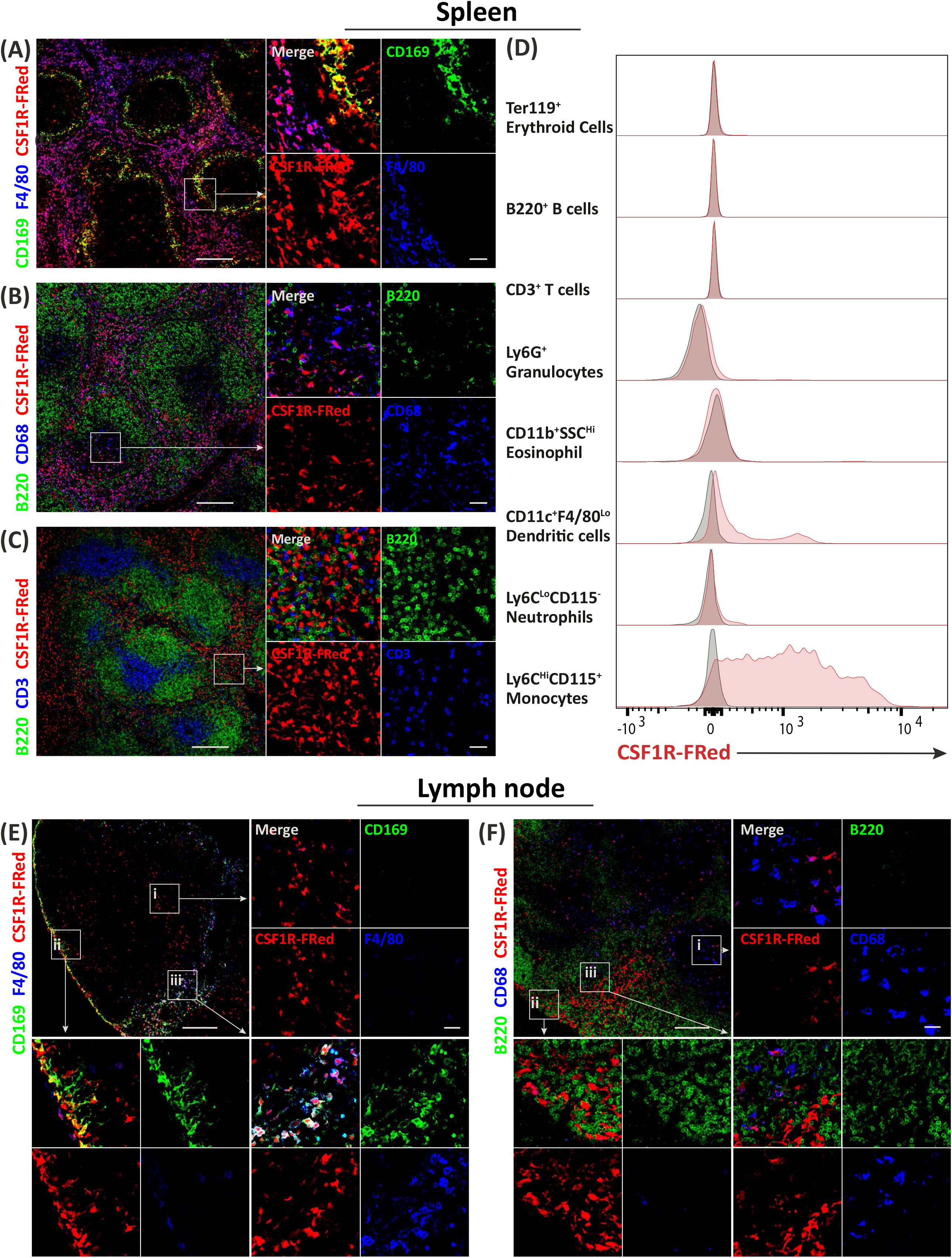
CSF1R-FRed expression in spleen and lymph node. Spleens and inguinal lymph nodes were harvested from adult wild-type and CSF1R-FRed +/- and fixed and sectioned as described in Materials and Methods. Part of the spleen was disaggregated for Flow Cytometry. CSF1R-FRed was detected with anti-RFP antibody in combination with the markers indicated. Panel A and insets highlight the expression of CSF1R-FRed in F4/80_+_ red pulp macrophages, F4/80_−_, CD169_+_ marginal zone metallophils and F4/80_−_, CD169_−_ marginal zone macrophages. Panel B highlights CSF1R-FRed_+_ cells within T cell areas (excluded B220) and the overlaps with CD68. Panel C and inserts show the exclusion of CSF1R-FRed from CD3_+_ T cells and B220_+_ B cells in the red pulp but the potential for interactions amongst the 3 cell types. Panel D shows representative FACS profiles for WT and CSF1R-FRed spleen (red histograms) separated based upon the gating strategy in **Figure S7.** Panel E shows identifies CSF1R-FRed expression in CD169_−_, F4/80_−_ interdigitating cells in T cell areas (nset (i)), CD169_+_, F4/80_−_ subcapsular sinus macrophages (inset (ii)), and CD169_+_, F4/80_+_ medullary sinus macrophages in the lymph node. Colocalisation in Panel F with B220 and CD68 further highlights interdigitating cells in T cell areas (inset (i)) and intimate interaction with B cells in the medullary cords/sinuses (insets (ii) and (iii)).

Like MZ metallophils in the spleen, the majority of subcapsular sinus macrophages (SSM) in LN express CD169 and lack F4/80 (63), although there is also an F4/80_+_ population on the LN surface (65). SSM depend upon CSF1 produced locally (63). There was almost complete overlap between CSF1R-FRed and CD169 in the subcapsular region, whereas F4/80 was detected on only a subset of cells. CSF1R-FRed was detected on the dense population of F4/80_hi_, CD169_+_ macrophages that populate the medullary sinuses (65) and was also detectable on the network of F4/80_−_/CD169_−_ interdigitating cells (66) found throughout T cell areas of the LN. These may include the population of T zone macrophages described by Baratin et al. (67) as well as cDC. In spleen and LN there were few positive cells in B cell areas and there was no apparent reactivity in the so-called tingible body macrophages.

The intestinal wall contains multiple macrophage and DC notably the CD169_hi_ populations surrounding the crypts. All of the lamina propria populations are acutely depleted by anti-CSF1R antibody, with consequent impacts on epithelial differentiation (68). **Figure S1** shows CSF1R-FRed expression in the small intestine. Expression was coincident with F4/80 in the stellate populations of the lamina propria, but expression of FRed also provided a unique marker for the abundant CSF1-dependent phagocyte population underlying the dome epithelium of the Peyer’s patch which lacks expression of markers such as F4/80, CD169, CD206 and CD64(69).

### CSF1R-FRed expression in known and novel resident tissue MPS cells

*Csf1r* reporter transgenes identified an extensive network of presumptive resident macrophages in every organ in the body (5, 6). **Figure 6** shows representative confocal images of multiple tissues from the adult CSF1R-FRed mouse highlighting the dense network of stellate interstitial cells. Direct whole mount imaging of fresh tissues without fixation highlights the full extent of the network of MPS cells. Detection of CSF1R-FRed provides improved visualisation of less-studied macrophage populations, including the extensive network of macrophages in skeletal and smooth muscle, and those of the uterus and cervix, stomach, adrenal gland, Islets of Langerhans, ovary, pancreas, brown fat and thymus.

**Figure 6.**
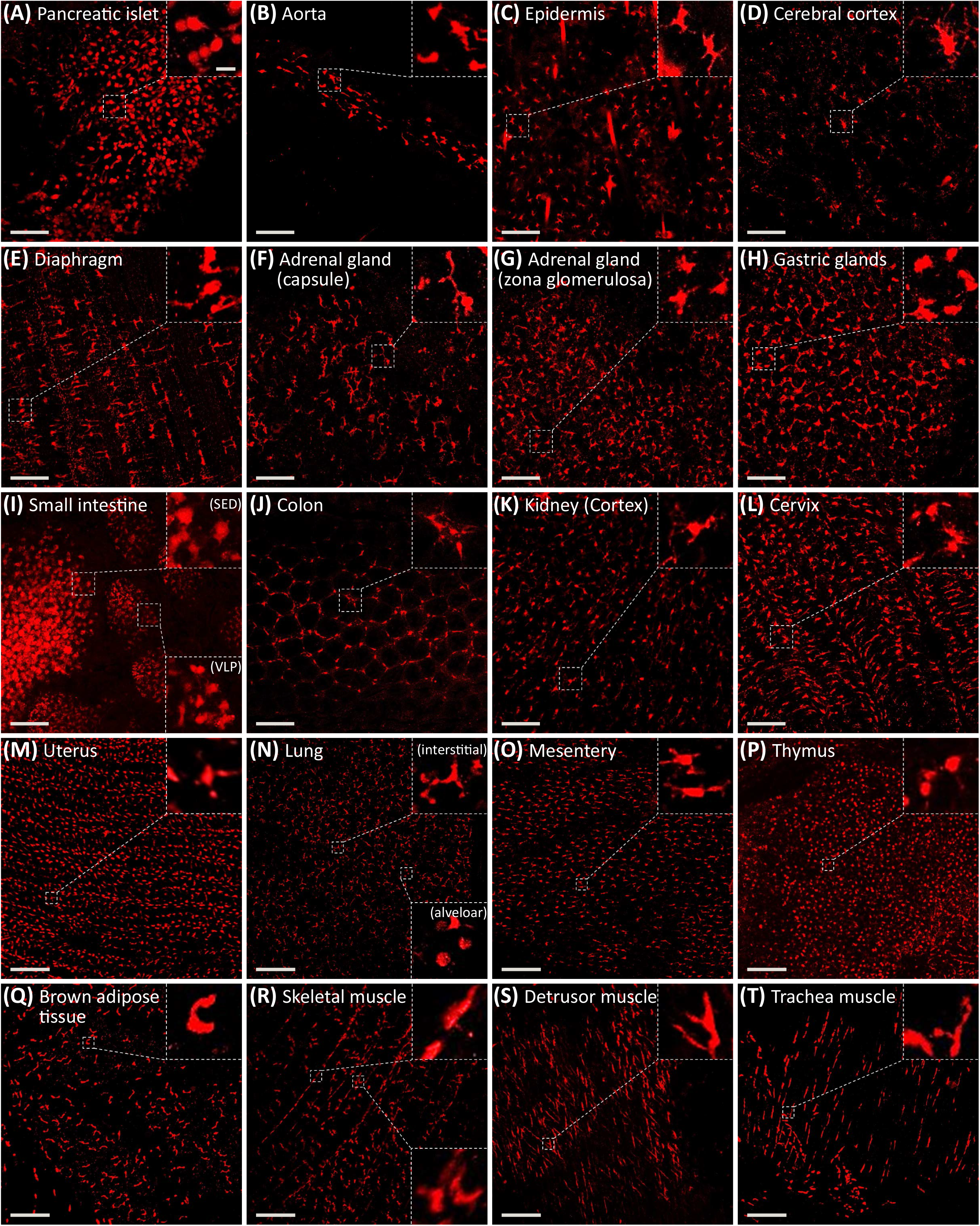
Imaging of macrophages in adult organs using CSF1R-Fred. Tissues were removed from adult CSF1R-FRed +/− mice and placed on ice in PBS. They were imaged directly within 2-3 hours using the Olympus FV3000 microscope. Images show maximum intensity projections of the tissues indicated. Many of these tissues can also be visualized as Z series in Supplementary **movies S1-S20**. Scale bar is 100μm.

As a multi-copy transgene, the *Csf1r*-EGFP reporter is considerably brighter than CSF1R-FRed. As discussed in the introduction, it has the disadvantage of expression in granulocytes and B cells, but it still has significant utility for live imaging. To determine the overlap between the two reporters, we crossed the CSF1R-FRed and *Csf1r*-EGFP lines. Recent studies have identified populations of CSF1R-dependent mononuclear phagocytes underlying the liver capsule and distinct from Kupffer cells (70, 71). These cells were implicated in defence against infiltration of the liver by pathogens in the peritoneal cavity. The liver capsular macrophages were detected using the expression of a *Cd207*-EGFP reporter transgene, a marker shared with Langerhans cells of the skin. The *Cd207*-EGFP reporter transgene was not detected in the capsule of other organs leading to the conclusion that the presence of these cells is unique to the liver (71). A confocal Z series inwards from the liver surface confirmed the detection of these cells with *Csf1r*-EGFP and CSF1R-FRed and the transition from the sub-capsular population to classical Kupffer cells **(Movie S1)**.

A similar series for the brain cortex (**Movie S2**) highlights the abundant macrophage population of the dura mater and leptomeninges(72) and the transition to the underlying network of microglia. Luo *et al.* (73) reported the expression of *Csf1r* in neurons and claimed that systemic CSF1 treatment could ameliorate neuronal injury by direct signalling. Contrary to that report, expression of the *Csf1r*-EGFP transgene is entirely restricted to microglia and brain-associated macrophages (20, 21). However, there is the formal possibility that the *Csf1r*- EGFP reporter construct lacks elements required for expression in neurons. **Figure S2** shows that CSF1R-FRed expression was not detectable in neurons in the dentate gyrus of the hippocampus, even in CSF1R-FRed_+/+_ mice with additional amplification provided by anti-RFP antibody.

The population of surface-associated or subcapsular macrophages was not restricted to the liver. The CSF1R reporter transgenes enable confocal imaging of the surface of every major organ of the body and reveal that surface-associated macrophages are a universal feature. The relative density and stellate morphology of surface-associated macrophages is remarkably consistent. In each case, a confocal Z series from the outer surface reveals transitions in morphology and location of the underlying resident tissue macrophages. **Movies S3-S17** show Z series for the two reporters through the depth of a selection of organs including the liver, lung, heart, brain, kidney, large intestine, periosteum, skeletal muscle, abdominal wall, epididymis, vas deferens adrenal and bladder. There was a complete overlap with the GFP and FRed reporter. Separate static and merged images of each reporter in selected tissues are shown in **Figure S3**. In each case, the Z series highlights the way in which the polarity of the macrophage spreading changes in various layers to align with the orientation of surface connective tissues/serosa/mesothelial layers, underlying muscle fibres, connective tissues and other structures.

To confirm the macrophage-restricted expression and utility for the isolation of other macrophage populations, we used conventional disaggregation procedures to isolate cells from liver, lung and brain and characterised the populations by Flow Cytometry **(Figure 7).** In each organ, expression was excluded from both lymphocytes and CD45-populations. In the liver, there is a complete overlap between CSF1R-FRed and F4/80 labelling of sinusoidal Kupffer cells **(Figure S4).** In isolated cells, the expression of CSF1R-Fred increased progressively from monocytes to mature TIM4_+_ Kupffer cells (**Figure 7A)** consistent with gene expression analysis of this progression (74). In the lung, CSF1R-FRed was expressed by interstitial monocyte and macrophage populations (**Figure 7C**). The expression in alveolar macrophages (F4/80_int_/CD11b_lo_) analysed by Flow Cytometry is obscured by very high autofluorescence and the relatively low *Csf1r* mRNA (2). Autofluorescence is less of an issue in confocal imaging. We confirmed expression in bronchial and alveolar populations and in interstitial and subcapsular cells *in situ* **(Figure 7B).** In the brain, CSF1R-FRed was detected in both microglia (CD45_lo_, CD11b_hi_) and brain-associated macrophages (CD45_hi_, CD11b_lo_) **(Figure 7D)** and consistent with **Figure S2** was absent from CD45-cells.

**Figure 7.**
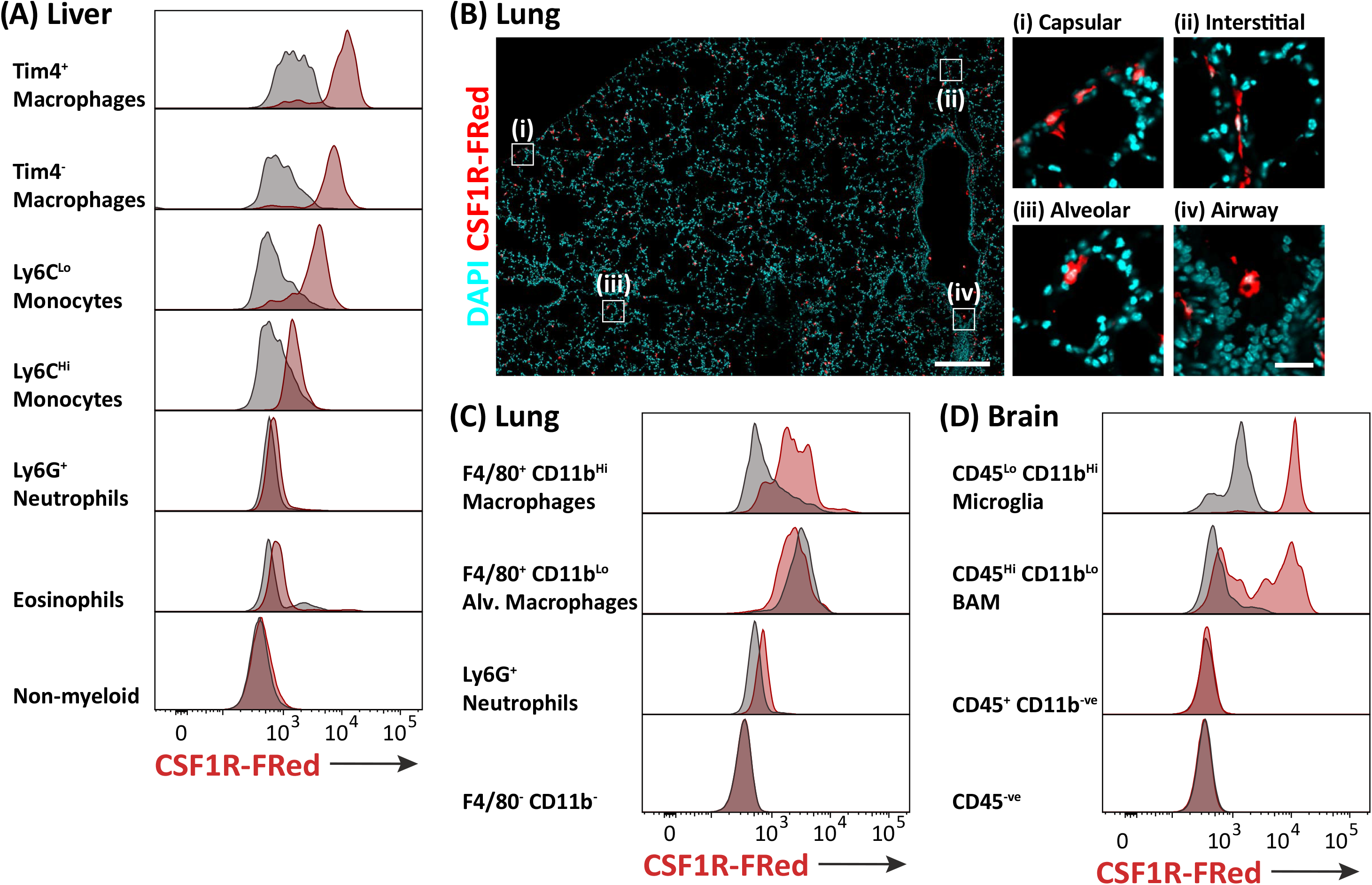
Expression of CSF1R-FRed in liver, lung and brain. Leukocytes were isolated from liver (A), lung (C) and brain (D) following tissue disaggregation as described in Materials and Methods and analysed by Flow Cytometry for expression of the indicated markers. The gating strategies are shown in **Figure S7.** Note in Panel (C) that alveolar macrophages have high levels of autofluorescence which may obscure a FRed signal. To confirm expression in lung macrophage populations, Panel B shows localization of CSF1R-FRed in unfixed sections lung. Inserts highlight four CSF1R-FRed_+_ populations distinguished by location and morphology. Scale bars

## Discussion

We have described the generation and characterization of a novel CSF1R reporter mouse line. The *Csf1r* promoter region including the conserved FIRE element has been used to drive reporter transgenes and to direct expression of constitutive and tamoxifen-inducible cre-recombinase and the macrophage Fas-induced Apoptosis (MaFIA) transgene for conditional macrophage ablation (3). These applications have been compromised by the expression of *Csf1r* mRNA and/or *Csf1r* promoter-driven transgenes in other hematopoietic cells and the potential for ectopic expression in multicopy transgenes. Many other myeloid promoters and target loci (e.g. *Adgre1, Ccr2, Cd68, Cd169, Cd11c, Cd11b, Clec9a, Cx3cr1, Flt3, LysM, Ms4a3, Tmem119, Zbtb46*) have been used to drive myeloid-restricted expression of reporters, cre-recombinase, diphtheria toxin receptor or other genes of interest. However, none of these alternatives is expressed universally and exclusively at the mRNA or protein level in all cells of the MPS (2). The CSF1R-FRed reporter mouse overcomes the confounding issues, providing a specific and robust MPS reporter for a large range of technical applications.

One future application of the *Csf1r-FRed* allele lies in study of the transcriptional regulation of the *Csf1r* locus. Deletion of the FIRE enhancer within intron 2 of *Csf1r* led to the loss of expression in monocytes and various tissue macrophages, including microglia, whereas other macrophage populations were unaffected. These impacts could only be assessed in homozygous mutant mice by the loss of the affected cell population (4). Several candidate enhancers other than FIRE have been identified in the *Csf1r* locus (4). Their role may be assessed by targeted deletion in CSF1R-FRed mice and analysis of expression of the mutated allele on a heterozygous *Csf1r* WT background.

From a purely technical viewpoint, the outcome establishes the feasibility of inserting any desired expression cassette (e.g. other reporters, gene of interest, Cre, DTR) into the mouse *Csf1r* locus to achieve MPS-restricted expression without disrupting expression or function of CSF1R protein. We have bred the *Csf1r-FRed* allele to homozygosity and there is no detectable impact on viability, growth, monocyte-macrophage abundance, surface markers (e.g. F4/80) or the expression of CSF1R on the cell surface **(Figure 3).** Accordingly, for many applications CSF1R-FRed can be maintained as a homozygous line and conveniently crossed to other reporters (as exemplified by the cross to the *Csf1r*-EGFP reporter). We note also that the FRed reporter is expressed in macrophages of homozygotes at around twice the level in heterozygotes increasing the signal and allowing convenient breeding **(Figures 3, 5)**. It is not self-evident that this would be the case, since the expression of *Csf1r* mRNA is regulated by CSF1R signals and the expression from the two alleles might be coordinated (75). The reciprocal observation is that in heterozygous *Csf1r* knockout mice and rats, there is no dosage compensation and both *Csf1r* mRNA and protein are reduced by 50% on monocytes and macrophages (1). It appears therefore that *Csf1r* alleles are regulated independently of each other.

The expression of CSF1R-FRed in BM accords with other evidence that *Csf1r* mRNA is absent from HSC and induced during lineage commitment downstream of expression of lineage-specific transcription factors (50, 53). The large majority of lineage-negative, CD115_+_ cells in BM are in cell cycle (25). The lack of detectable CSF1R-FRed in HSC is not compatible with the proposal that CSF1 acts directly on HSC to instruct myeloid lineage commitment via induction of PU.1 (52). These studies relied on detection of *Csf1r* in HSC by qRT-PCR and did not detect the CSF1R protein. The reported effects of CSF1 on PU.1 expression in isolated HSC were small and significant numbers of PU.1-expressing cells arose spontaneously even in the absence of added CSF1. Accordingly, we favour a selective model of CSF1 action. Interestingly, although CSF1 administration can induce a substantial monocytosis, CSF1R signaling is not absolutely required for monocyte production. Anti-CSF1R antibody treatment of mice depleted tissue macrophages but had no effect on the blood monocyte count (23, 24) and mutation of the *Csf1r* FIRE enhancer ablated expression in BM and blood monocytes without impacting abundance or phenotype of these cells (4). CSF1R-FRed was more highly-expressed and somewhat heterogeneous within CMP compared to GMP as defined using CD32 as a marker (55). Several recent studies question the hierarchical model of myelopoiesis in which CMP give rise to GMP. They indicate that CD115hi monocyte-DC progenitors (MDP) (32) and committed monocyte progenitors (25) may both reside within the CMP fraction of BM ((76) and references therein). Conversely, Kwok et al. (77) dissected the GMP populations to reveal a predominance of CD115_lo_ committed neutrophil progenitors consistent with low expression of CSF1R-FRed in this fraction **(Figure 3B)** and the fact that granulocyte express *Csf1r* mRNA but do not translate the protein (12).

One novel finding in the BM was the expression of CSF1R in megakaryocytes (**Figure 3D**). Expression profiling of mouse megakaryocyte differentiation *in vitro* revealed high expression of *Csf1r* mRNA which declined in the most mature population (78). This conclusion is consistent with detection of CSF1R-FRed in the MEP population **(Figure 3C).** The functional importance of CSF1/CSF1R in megakaryocyte biology is unknown. In humans, *CSF1R* is commonly deleted in 5q-syndrome, which is associated with thrombocytopenia (79) but homozygous mutation in *CSF1R* in rats or humans had no impact on platelet count (80, 81). Conversely, in humans, mice and pigs, CSF1 administration causes a rapid fall in platelet count (82–85). In mice, this drop resolved with a significant overshoot, even with continued CSF1 treatment, associated with alterations in megakaryocyte ploidy (83). CSF1-induced thrombocytopenia was attributed to reduced circulating half-life; the possibility that CSF1 might also drive the rebound was not excluded.

Classical DC in spleen, LN and also in Peyer’s patches form a dense network of interdigitating cells within the T cell areas (66). The *Csf1r*-EGFP transgene was expressed by interdigitating cells in spleen and LN T cells areas and by CD11c_hi_ DC isolated from spleen and lymph node and surface CD115 was detected on a subset of these cells (35). The level of *Csf1r* mRNA also distinguishes CD8_+_ (cDC1) and CD8-DC isolated from spleen (BioGPS.org, Immgen.org) and cDC1 (XCR1_+_, CD103_+_) from cDC2 (CD11b_+_) in multiple large RNA-seq datasets (2), with relatively higher expression by cDC2. CSF1R-FRed was detected at varying levels in isolated CD11c_hi_ spleen DC and may provide a subset marker **(Figure 6).** Given the extensive ramification of the interdigitating cells within T cell areas, it is likely that they are under-represented in populations obtained by enzymic disaggregation. The expression of CSF1R-FRed is consistent with evidence that cDC in lymphoid tissues can be regulated by CSF1R signals (34, 35).

Other macrophage populations in spleen and LN that have not been isolated by tissue disaggregation include those of the MZ and subcapsular sinus (SCS). We co-localised CSF1R-FRed with F4/80 (which is excluded from the marginal zone and T cell areas) and CD169 (expressed by both MZ and SCS macrophages) **(Figure 5)**. CSF1R-FRed was detected in red pulp (F4/80_hi_) and marginal zone (CD169_+_, F4/80_−_) macrophages. Unexpectedly, CSF1R-FRed was undetectable in the large tingible-body macrophages in germinal centres which do express *Csf1r*-EGFP but are CSF1R-independent (23). These cells express CD68 and MERTK, which is required for apoptotic B cell uptake (86). Unlike follicular DC they are BM-derived (87) but they also lack other myeloid markers including IBA1 and F4/80 (88, 89).

Flow cytometry analysis and imaging of multiple tissues confirms that, like *Csf1r* mRNA, CSF1R-FRed is expressed throughout the MPS. By contrast to previous reports (73, 90, 91) we found no evidence of CSF1R protein expression in brain neuronal cells or epithelial cell populations in intestine or kidney. Accordingly, the numerous pleotropic impacts of *CSF1R* mutations and kinase inhibitors in humans and experimental animals can be attributed entirely to indirect consequences of MPS cell deficiency (reviewed in (1).

In non-lymphoid tissues, there was a complete overlap of CSF1R-FRed with *Csf1r*-EGFP, supporting the utility of this transgenic marker **(Figure S3).** Not surprisingly, as a multi-copy transgene the signal from *Csf1r*-EGFP is generally much brighter than CSF1R-FRed, but the signal intensities are not perfectly correlated. There are cells in several of the tissues analyzed that appear more red than green, perhaps reflecting the fact that the transgenic reporter does not contain all the *Csf1r* regulatory elements (4). Both transgenes viewed in unfixed fresh tissues highlight the full extent of the MPS network in every tissue. *Csf1r* mRNA is readily detected in total mRNA from all the organs shown at levels around 10% of that detected in BMDM (92) so the density of these cells is not surprising. Within tissues, CSF1R-FRed_+_/*Csf1r*-EGFP_+_ cells appear spread on surfaces such as epithelial and endothelial basement membranes and muscle and connective tissue fibres. Both reporters highlighted the so-called tenophages recently identified in tendon (93). The localization of CSF1R-FRed/*Csf1r*-EGFP confirmed the existence of surface-associated macrophages underlying the serosa and spread in the plane of the capsule in every major organ. They include the heterogeneous population of macrophages associated with the dura mater of the brain, previously characterized using CX3R1, MHCII, CD169, IBA1 and CD11c (72) and recently isolated and profiled as a distinct population from microglia (94). Conversely, the dense subcapsular macrophage population of the lung was not recognized or characterized in studies of interstitial macrophage heterogeneity (95). These surface-associated macrophage populations present in every organ were identified in the original studies of the localization of the F4/80 antigen (reviewed in (15)) but are not readily visualized in cross section. It remains to be determined whether they have organ-specific innate immune or homeostatic functions or share phenotypes with the cells identified in the liver (70, 71).

One location where the abundance of resident macrophages has been under-appreciated is muscle. The macrophages of the smooth muscle of the intestinal muscularis have been attributed functions in interactions with enteric neurons and the control of peristalsis and gut motility (96). Whole mount imaging of these cells with the CSF1R-FRed and *Csf1r*-EGFP reporter reveals their abundance within the myenteric plexus, the way in which they spread along the muscle fibres, and change polarity between the longitudinal and circular muscle layers. The same transitions are evident in layers of smooth muscle in large intestine, abdominal wall, diaphragm, uterus/cervix, vas deferens and bladder as well as cardiac muscle (**Figures 6A, 6E, 6L, 6M; Movies S4, S8, S9, S12, S17**). By contrast, there is very limited literature on macrophages in skeletal muscle, where CSF1R-FRed-positive cells are at least as abundant as in smooth muscle, with the same regular spacing along the muscle fibres (see **Figures 6R-T; Movie S7**). Previous studies using immunohistochemical markers (CD11b, F4/80, CD45) identified abundant resident macrophage-like populations in the epimysium and perimysium (the external muscle envelope), but rarely detected positive cells in the endomysium (the thin layer between muscle fibres) (97). We suggest this is a reflection of their extensive spreading, and detection further compromised by fixation (which causes the macrophages to contract) and sectioning (which tears them from the muscle fibres). Standard methods of tissue disaggregation yield very few macrophage-like cells from undamaged muscle. The resident tissue macrophages in muscle are well-positioned to interact with satellite cells and the neuromuscular junctions and to contribute similar regulatory, homeostatic and remodelling functions to resident macrophages identified in the heart and vascular smooth muscle (98–100) that were also readily visualized with CSF1R-FRed and *Csf1r*-EGFP.

The locations occupied by tissue macrophages have been referred to as niches or territories. The concepts differ depending on whether the precise location is physically defined (a niche) or macrophages define their own territory by mutual repulsion (16, 101). The regular spacing in every location **(Figure 6)** supports the territory concept. The two concepts are compatible if a niche exists on a macroscopic scale (e.g. a particular surface) and macrophages determine their density within the niche. Almost universal expression of semaphorins and their receptors (known regulators of cell motility) (2) in isolated tissue macrophages from all organs provides at least one possible mechanism for communication between resident macrophages.

In summary, we have produced a novel transgenic line that provides a specific and ubiquitous marker for MPS cells and highlights their distribution in every organ of the body. Given the abundance of these cells, it is not surprising that their depletion in CSF1R-deficient mice, rats and humans has profound impacts on postnatal growth and development (1).

## Materials and Methods

### Generation of Csf1r-T2A-FusionRed mice

We used CRISPR-Cas9 to generate CSF1R C-terminally tagged with a T2A-FusionRed reporter gene. The overall strategy is shown in **Figure 1.** Two sgRNA targeting the STOP codon of *Csf1r* were selected using the Sanger WTSI website (http://www.sanger.ac.uk/htgt/wge/). Guides present elsewhere in the genome with mismatch (MM) of 0, 1 or 2 were discounted, MM3 were considered if predicted off targets were not exonic. sgRNA sequences (aactaccagttctgctgaag-*tgg* and actaccagttctgctgaagt-*ggg*) were purchased as crRNA oligos, which were annealed with tracrRNA (both oligos supplied IDT; Coralville, USA) in sterile, RNase free injection buffer (TrisHCl 1 mM, pH 7.5, EDTA 0.1 mM). 2.5 μg crRNA was combined with 5 μg tracrRNA, heated to 95 °C, then allowed to slowly cool to RT.

For the donor repair template we used the EASI-CRISPR long-ssDNA strategy (102) which comprised the FusionRed gene with cleavable T2A linker flanked by 136 nt and 139 nt homology arms. The lssDNA donor was generated using Biotinylated PCR from a dsDNA template, followed by binding to Streptavidin beads and on-bead denaturation to remove the bottom strand of DNA. The top, single stranded DNA was then cleaved off the column by hybridizing a short oligo at the 5’ end to reform a *KpnI* restriction site and subjected to restriction digestion.

For embryo microinjection the annealed sgRNA was complexed with Cas9 protein (New England Biolabs) at RT for 10 min, before addition of long ssDNA donor (final concentrations; sgRNA 20 ng/μL, Cas9 protein 20 ng/μL, lssDNA 10 ng/μL). CRISPR reagents were directly microinjected into C57BL/6JOlaHsd zygote pronuclei using standard methods (103). Zygotes were cultured overnight and the resulting 2 cell embryos surgically implanted into the oviduct of day 0.5 post-coitum pseudopregnant mice. Potential founder mice were screened by PCR, first using primers that flank the homology arms and sgRNA sites (Geno F2 ccaccccaggactatgctaa and Geno R2 ctagcactgtgagaacccca), a reaction which both amplifies the WT band (394 bp), any InDels that result from NHEJ activity and larger products (1153 bp) indicating HDR **(Figure 1).** Pups with a larger band were reserved, the band isolated and amplified again using high fidelity Phusion polymerase (NEB), gel extracted and subcloned into pCRblunt (Invitrogen). Colonies were mini-prepped and Sanger sequenced with M13 Forward and Reverse primers. Pups showed perfect sequence integration after alignment of sequence traces to predicted knock-in sequence and bred with a WT C57BL/6JOlaHsd to confirm germline transmission and establish the colony. The founders were subsequently transferred to SPF facilities in Edinburgh and Brisbane by rederivation and crossed to the C57BL/6JCrl background.

### Tissue collection for imaging and disaggregation for flow cytometry analysis

Peripheral blood (100 μL) was routinely collected into EDTA tubes by cardiac puncture following euthanasia. Blood was subjected to red blood cell lysis for 2 min in ACK lysis buffer (150 mM NH_4_Cl, 10 mM KHCO_3_, 0.1 mM EDTA, pH 7.4) and resuspended in flow cytometry (FC) buffer (PBS/2 % FBS) for staining. Peritoneal cells were recovered by lavage with 10 mL PBS. Following lavage tissues of interest were removed for disaggregation and imaging. Tissues for imaging were stored in PBS on ice and imaged within 2 h.

Liver and brain tissues for disaggregation were finely chopped in digestion solution containing 1 mg/mL Collagenase IV, 0.5 mg/mL Dispase (Worthington) 20 μg/mL DNAse1 (Roche) and placed on ice until further processing (~1 g tissue/10 mL). Tissues in digestion solution were placed on a rocking platform at 37°C for 45 min prior to mashing through a 70 μm filter (Falcon). For liver and brain Percoll density gradient centrifugation was used to isolate the non-parenchymal fraction, as previously described (4, 5). Cell isolation from BM, spleen and lymph node (LN) was carried out as described (54). 5 × 10_6_ cells were stained for flow cytometry analysis and 1-2 × 10_6_ cells were acquired for analysis.

### Flow cytometry

Cell preparations were stained for 45 min on ice in 2.4.G2 hybridoma supernatant to block Fc receptor binding. Myeloid populations were stained using antibody cocktails (54) comprising combinations of CD45-BV421, F4/80-AF647, Cd11b-BV510, Cd115-BV605, Ly6G-BV785 or APC-Fire750, Tim4-PECy7, Ly6C-PB or BV785, B220-APCCy7 or BUV496, TER119-BUV395 and CD3-PE-Cy5 (Biolegend). Hematopoietic stem and progenitor cells were stained using a standard antibody cocktail comprising of Lineage-BV785 (CD3-, CD45R-, CD41-, CD11b-, GR1- and TER119-biotin followed by streptavidin-BV785), cKIT-APC, SCA-1-PE-Cy7, CD150-BV650, CD48-BV421 and CD115-BV605 (Biolegend). Myeloid and lymphoid progenitors were stained for using an antibody cocktail comprising of Lineage-BV785 (CD3-, CD45R-, CD11b-, GR1- and TER119-biotin followed by streptavidin-BV785), c-KIT-APC, SCA-1-PE-Cy7, CD34-PerCP-Cy5.5, CD16/32-BV421, FLT3-APC-Cy7. Cells were washed twice following staining and resuspended in FC buffer containing 7AAD (Life Technologies) or FVS700 (BD Bioscience) for acquisition using a Cytoflex (Beckman Coulter) or Fortessa (Becton Dickinson). Relevant single-color controls were used for compensation and unstained and fluorescence-minus-one controls were used to confirm gating strategies. Flow cytometry data were analyzed using FlowJo 10 (Tree Star). Live single cells were identified for phenotypic analysis by excluding doublets (FSC-A > FSC-H), 7AAD_+_ or FVS700_+_ dead cells and debris. Gating strategies for each of the analyses in this study are shown in **Figures S5, S6** and **S7**.

### Confocal microscopy and immunofluorescence

All immunofluorescent images were acquired on the Olympus FV3000 microscope (Olympus Corporation) and FLUOVIEW software (Olympus Corporation). To prepare tissues for immunofluorescent staining: spleens, LN, livers and lungs were snap frozen in liquid nitrogen, and bones were first fixed in 4% paraformaldehyde. Bones were then further decalcified in 14% EDTA over 3 weeks with several changes before sucrose saturation in 15-30% sucrose gradient over 2 days. All tissues were then embedded in Optimal cutting temperature compound (ProSciTech) and cut at 5 μm thickness. FusionRed in the bones was stained for using Rabbit anti-RFP antibody (Abcam ab62341). Cell specific staining for macrophage phenotypic markers was performed using a monoclonal Rat anti-F4/80 (Abcam) or directly conjugated anti-CD169-AF488 or anti-CD68-AF647 (both Biolegend). B cells and T cells were detected using anti-B220-AF488 (Biolegend) and anti-CD3-AF647 respectively (BD Bioscience). Unconjugated primary antibodies were detected with fluorescently labeled species appropriate secondary antibodies including Goat Anti-Rabbit IgG (H+L) Alexa Fluor 594 (ThermoFisher Scientific) and Goat Anti-Rat IgG (H+L)-Alexa Fluor 647 (ThermoFisher Scientific). All sections of bone, spleen, LN, liver, lungs and brain images were acquired with a 200x magnification objective using the multi-area tile scan function. For direct imaging of wholemount tissues, tissues were left as undisturbed as possible for direct imaging through the tissue surface. For tissues with reduced laser penetration (such as spleen or kidney) tissues were trimmed to a relative thickness of 2 mm and imaged from the surface in. The Z-stack function of the software was used to acquire a greater depth of field. For wholemount imaging of small intestine, 1 mm pieces of small intestine were fixed for 1 h in BD Cytofix/Cytoperm buffer before staining with a rat anti-F4/80 monoclonal (Abcam). Secondary staining was performed using an anti-rat AF647 (ThermoFisher Scientific) and B cells were stained using anti-B220-AF488 (Biolegend). CSF1R-FRed signal was acquired using the 561-diode laser and *Csf1r*-EGFP or AF488 was acquired using the 488-diode laser both on the FV3000 main combiner (FV31-MCOMB). For DAPI or AF647, signal was acquired using the 405- and 633-diode lasers respectively on the FV3000 main combiner (FV31-MCOMB).

## Supporting information

Movie S1 Liver

Movie S2 brain

Movie S3 heart

Movie S4 Colon

Movie S5 Lung

Movie S6 Kidney

Movie S7 muscle

Movie S8 Abdominal wall

Movie S9 Abdominal wall

Movie S10 tendon

Movie S11 pleural wall-rib

Movie S12 Bladdeer

Movie S13 Adrenal

Movie S14 Eye

Movie S15 Seminal vesicle

Movie S16 Epididymis

Movie S17 van deferens

## Acknowledgments

This work was supported by the Medical Research Council (MRC) UK grant MR/M019969/1 and by Australian National Health and Medical Research Council (NHMRC) grant GNT1163981 to DAH. DAH receives core support from The Mater Foundation. RR was supported by a doctoral scholarship (application number: 314413, file number: 218819) granted by CONACyT Nuevo Leon—I2T2, Mexico. PH was supported by the biotechnology and biological sciences research council (BBSRC) grant BB/P013732/1. DDO was supported by MRC grant MR/M010341/1. We acknowledge support from the Microscopy and Cytometry facilities of the Translational Research Institute (TRI). TRI is supported by the Australian Government.

## Supplementary Materials

**Figure S1.** Colocalization of CSF1R-Fred and markers of macrophages, B cells and T cells in small intestine and Peyer’s patch.

**Figure S2.** Colocalization of CSF1R-FRed and neuronal markers (NeuN, Tubulin 3) in hippocampus

**Figure S3.** Comparative localization of EGFP and FRed in CSF1R-FRed/Csf1r-EGFP mice.

**Figure S4.** Colocalization of CSF1R-FRed and F4/80 in liver.

**Figure S5.** Gating strategies for Flow Cytometry analysis of stem and multipotent progenitors in bone marrow

**Figure S6.** Gating strategies for Flow Cytometry analysis of committed progenitors in bone marrow.

**Figure S7.** Gating strategies for Flow Cytometry analysis of blood, peritoneum, spleen, liver, lung and brain.

## Supplementary movie descriptions

All movies are Z series of fresh tissue from CSF1R-FRed x *Csf1r*-EGFP mice without fixation. Each movie involves 1μm steps from the surface and the depth is shown.

**Movie S1. Liver.** Shows the transition from ramified subcapsular macrophage populations spread in the plane of the surface (0-25μm) to underlying Kupffer cells.

**Movie S2. Brain cortex.** Shows the transition from meningeal macrophages spread in the plane of the surface (0-15μm) to underlying microglia (15-37μm). Includes 3D reconstruction of microglial processes

**Movie S3. Heart.** Shows transition from epicardial macrophage populations spread in the plane of the surface (0-20μm) to macrophages spread along muscle fibres in the myocardium.

**Movie S4. Colon.** Shows transition from macrophages spread on the serosal surface (0-35μm) to macrophages spread long longitudinal and circular muscle fibres (35-70μm) to macrophages surrounding villi in the submucosa and mucosa.

**Movie S5. Lung.** Shows transition from subcapsular macrophages on the pleural surface (0-20μm) to underlying alveolar and interstitial macrophage populations.

**Movie S6. Kidney.** Shows transition from subcapsular macrophages spread in the plane of the surface to underlying interstitial macrophages surrounding tubules in the cortex (note that there is red autofluorescence in the tubules).

**Movie S7. Skeletal muscle (zygomaticus major).** Oblique series distinct morphologies of macrophages of the epimysium and transition to elongated macrophages spread along muscle fibres.

**Movie S8. Abdominal wall-peritoneal side.** Shows change in polarisation of macrophage populations aligned with changes in muscle fibre orientation.

**Move S9. Abdominal wall-skin side.** Shows change in polarisation of macrophage populations aligned with changes in muscle fibre orientation.

**Movie S10. Tendon (zygomaticus major).** Shows transition from macrophages on the surface to tendon macrophages spread longitudinally along connective tissue fibres.

**Movie S11. Pleural wall-ribs.** Shows macrophages aligned along intercostal muscle fibres and transition (10-45μm) to dense periosteal macrophage population on the surface of ribs.

**Movie S12. Bladder-detrusor muscle.** Shows macrophages spread longitudinally along muscle fibres at all depths (note muscle is spontaneously contracting).

**Movie S13. Adrenal gland.** Shows transition from subcapsular macrophages to populations of the zona glomerulosa. Note the tissue surface is not flat and subcapsular populations are progressively visualized on the edge of the field.

**Movie S14. Eye. Sclera and choroid.** Shows transition from the sclera (0-25μm) to the choroid layers of the eye. Note that the tissue is not flat and scleral populations are progressively visualized on the edge of the field.

**Movie S15. Seminal vesicle.** Shows the transition from a subcapsular macrophage population (0-20μm) through a fibromuscular layer to an underlying glandular structure region (45μm-80μm) where macrophages outline the surface of glands and ducts.

**Movie S16. Epididymis.** Shows the transition from a subcapsular macrophage population (0-20μm) to dense interstitial macrophage populations aligned with the surface of seminiferous tubules.

**Movie S17. Van deferens.** Shows the transition from a serosal/subcapsular macrophage populations of the outer layer (0-20μm) to populations aligned with the middle (longitudinal) muscle layer (20-40μm) and then at right angles, to the circular muscle layer (note muscle is spontaneously contracting).

