## Supplementary Figures S1-S7 for "A transgenic line that reports CSF1R protein expression provides a definitive marker for the mouse mononuclear phagocyte system"

### Supplementary Figure Legends

**Figure S1.** Wholemount imaging of CSF1R-FRed (red) Peyer's patches and villi of small intestine with co-immunostaining with F4/80 (blue) and B220 (green). SED, subepithelial dome; VLP, villus lamina propria. Main image scale bar = 500µm. Magnified images scale bars: (ii) = 50um; (iii) and (iv) = 20um.

**Figure S2.** Colocalization of CSF1R-FRed and neuronal markers (NeuN, Tubulin 3) in hippocampus.

**Figure S3.** Comparative localization of EGFP and CSF1R-FRed in CSF1R-FRed/Csf1r-EGFP mice. Main image scale bar = 100µm. Magnified images scale bars = 10µm.

**Figure S4.** Comparative localization of CSF1R-FRed (red) and F4/80 (blue) in the liver. Portal vein (PV), portal triad (PT) and central vein (CV) have been highlighted in magnified images. Open arrows indicate CSF1R-FRed$^{+}$F4/80$^{+}$ monocytes in the portal vein. Main image scale bar = 200µm. Magnified images scale bars = 20µm.

**Figure S5.** Gating strategies for Flow Cytometry analysis of (A) hematopoietic stem and progenitor cells, (B) common myeloid progenitors, (C) common lymphoid progenitors and (D) mature lineages in bone marrow.

**Figure S6.** Flow cytometry gating strategies for (A) peripheral blood, (B) peritoneal fluid, (C) spleen, (D) liver, (E) brain and (F) lung.

**Figure S7.** (A) RFP antibody isotype (Rabbit IgG) in CSF1R-FRed bones marrow and (B) anti-RFP immunostaining in bone marrow of WT (C57BL/6) bone marrow. Additional staining for CD169 (green) and F4/80 (blue) to identify bone marrow populations was also performed. Counterstained with DAPI. BV, blood vessel; MGK, megakaryocyte; OC, osteoclast. Main image scale bar = 200µm. Magnified images scale bars = 20µm.

**Supplementary movie descriptions.**

All movies are Z series of fresh tissue from CSF1R-FRed x *Csf1r*-EGFP mice without fixation. Each movie involves scanning in 1µm steps from the surface into the tissue and the depth is shown. Variation in depth scanned was dependent on confocal laser penetration into tissues. All scale bars = 100µm

**Movie S16. Epididymis.** Shows the transition from a subcapsular macrophage population (0-20µm) to dense interstitial macrophage populations aligned with the surface of seminiferous tubules.

**Movie S17. Van deferens.** Shows the transition from a serosal/subcapsular macrophage populations of the outer layer (0-20µm) to populations aligned with the middle (longitudinal) muscle layer (20-40µm) and then at right angles, to the circular muscle layer (note muscle is spontaneously contracting).

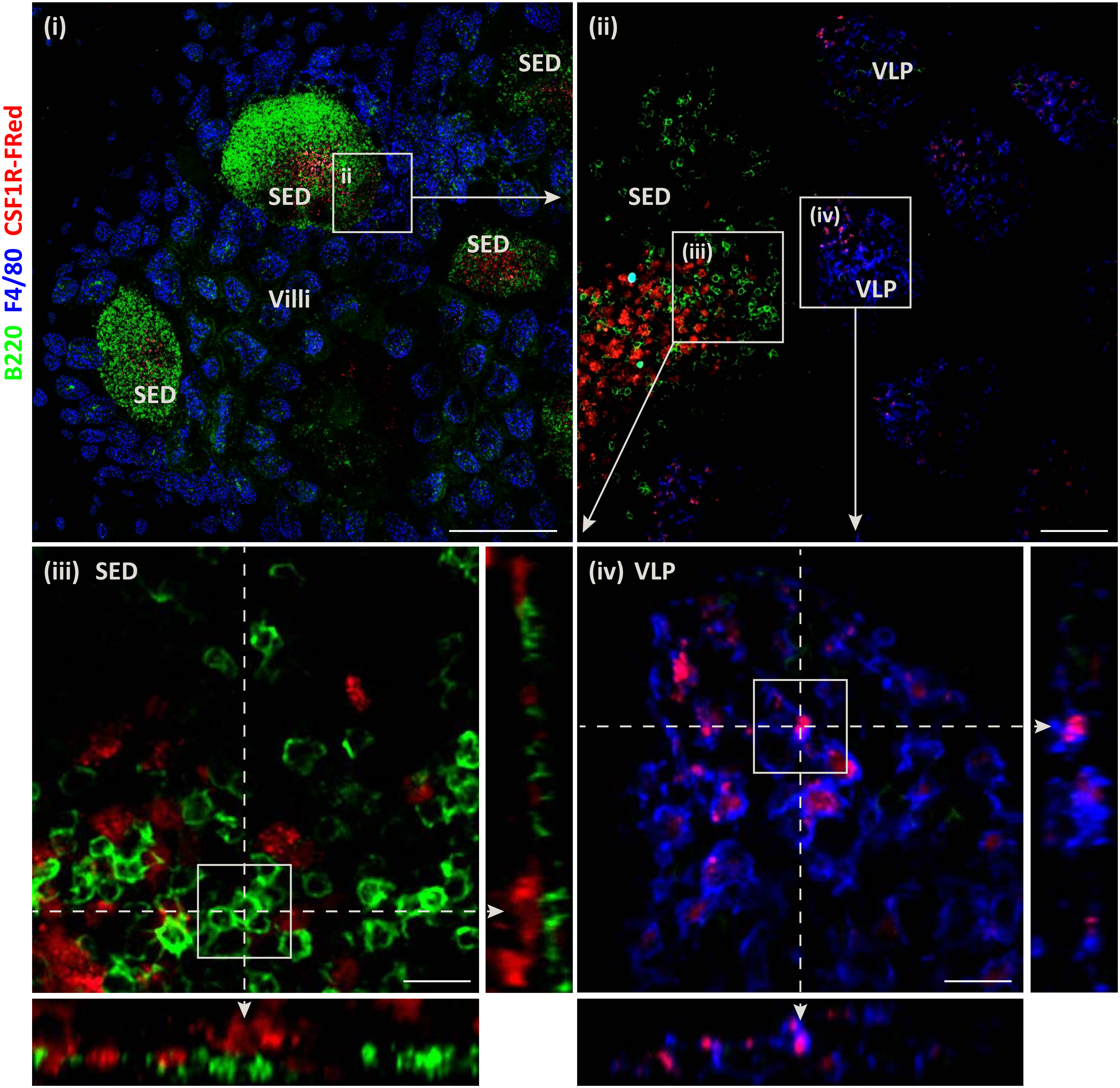
B220 F4/80 CSF1R-FRed
(i)
SED
ii
SED
SED
Villi
SED
(ii)
VLP
SED
(iv)
(iii)
VLP
(iii) SED
(iv) VLP

Figure S1

Figure S2A
Expression of CSF1R-FRed in the dentate gyrus of the hippocampus.
NeuN
CSF1R-FRed (Anti-RFP)
DAPI

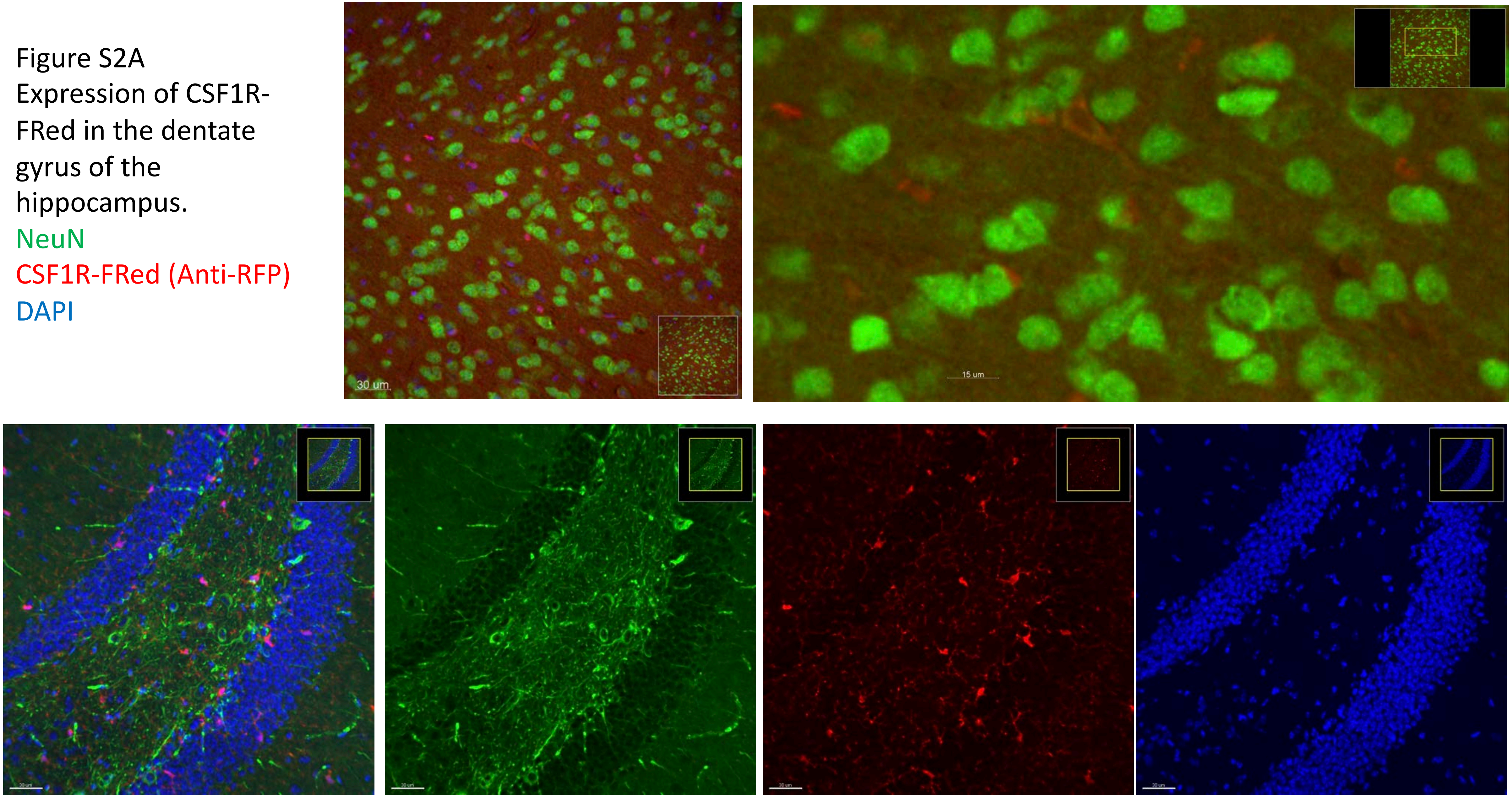

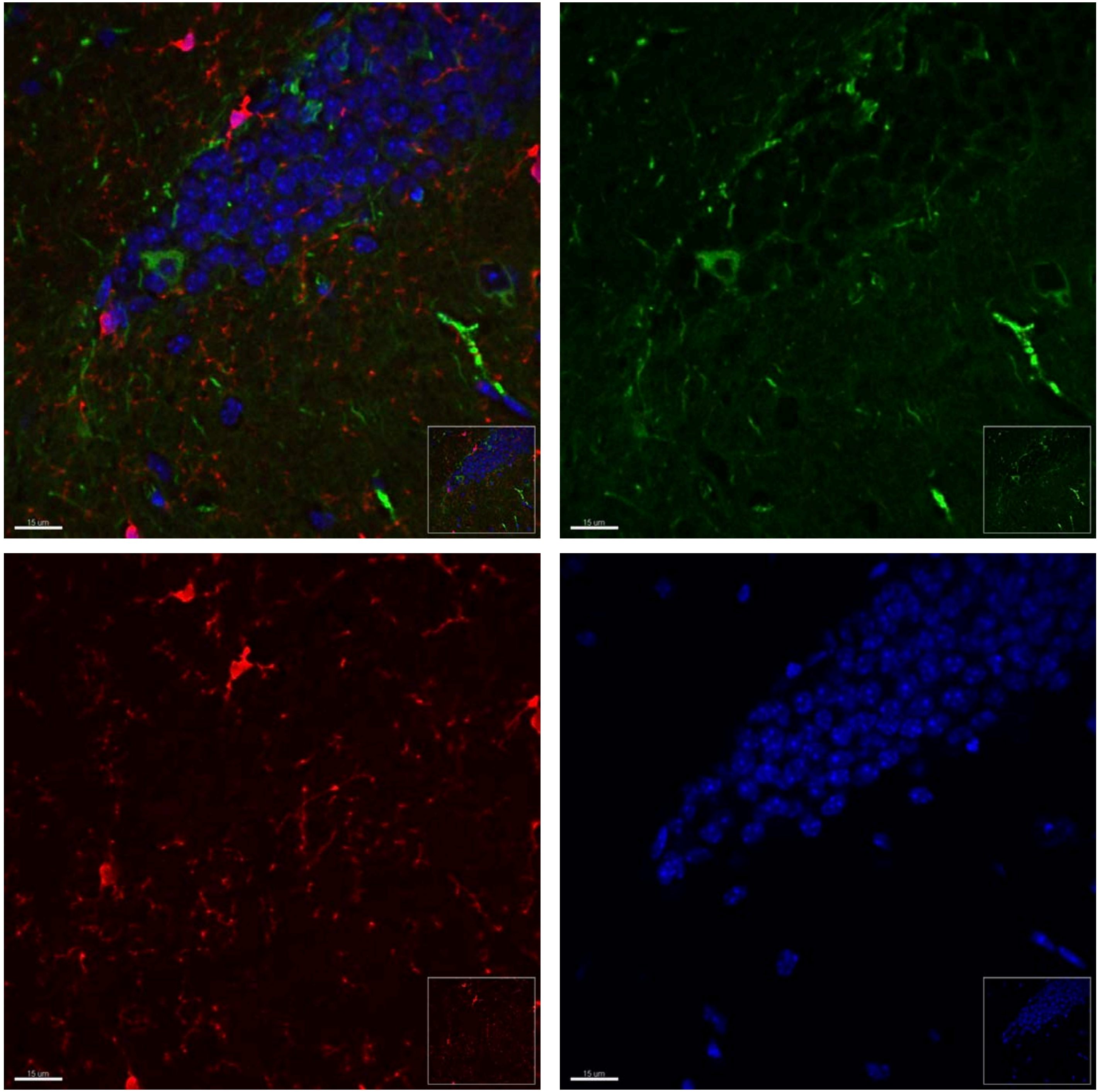


Figure S2B
Expression CSF1R-FRed in the dentate gyrus of the hippocampus.
(scale bar= 30$\mu m$).
Tubulin 3
CSF1R-FRed (Anti-RFP)
DAPI

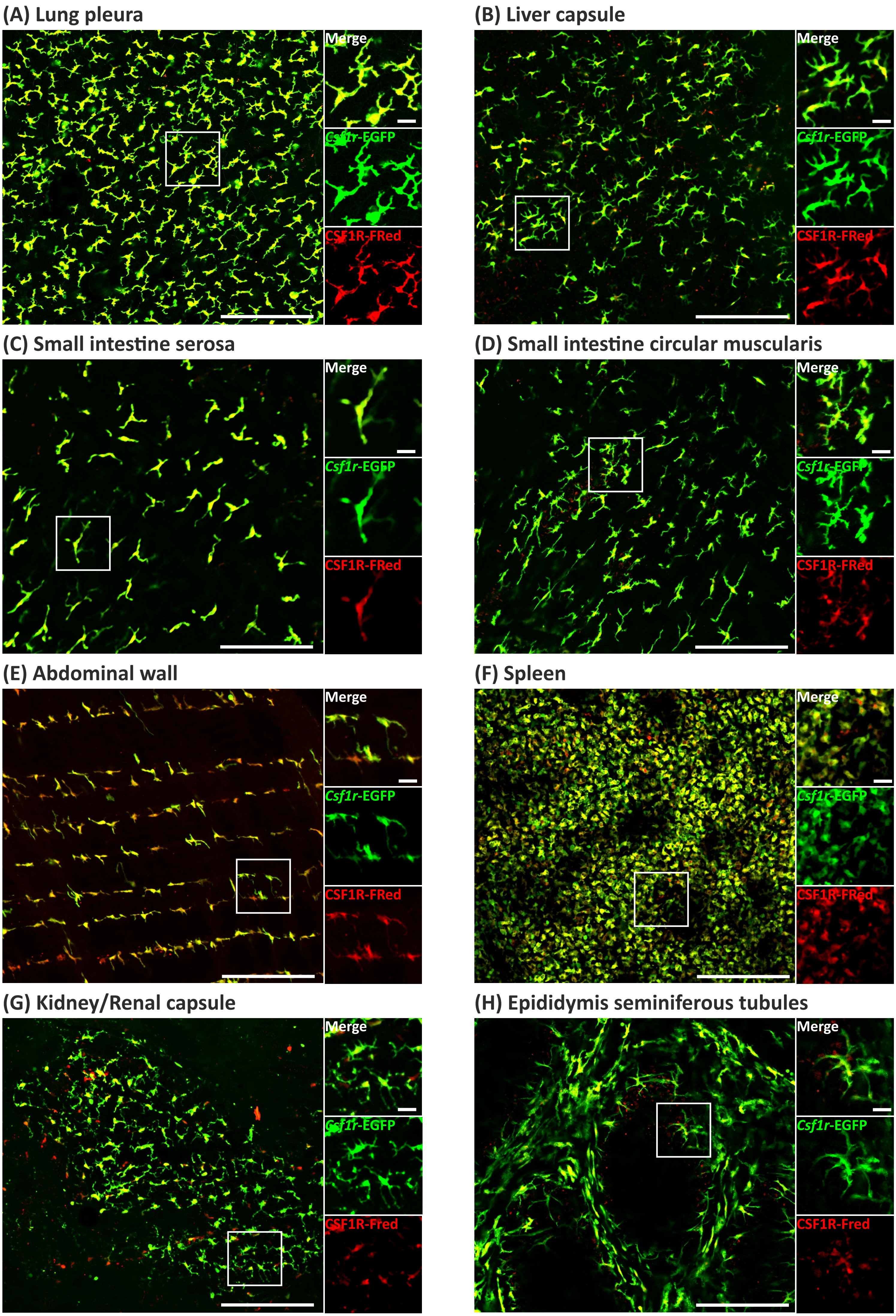
(A) Lung pleura
Merge
Csf1r-EGFP
CSF1R-FRed
(B) Liver capsule
Merge
Csf1r-EGFP
CSF1R-FRed
(C) Small intestine serosa
Merge
Csf1r-EGFP
CSF1R-FRed
(D) Small intestine circular muscularis
Merge
Csf1r-EGFP
CSF1R-FRed
(E) Abdominal wall
Merge
Csf1r-EGFP
CSF1R-FRed
(F) Spleen
Merge
Csf1r-EGFP
CSF1R-FRed
(G) Kidney/Renal capsule
Merge
Csf1r-EGFP
CSF1R-Fred
(H) Epididymis seminiferous tubules
Merge
Csf1r-EGFP
CSF1R-Fred

**Figure S3**

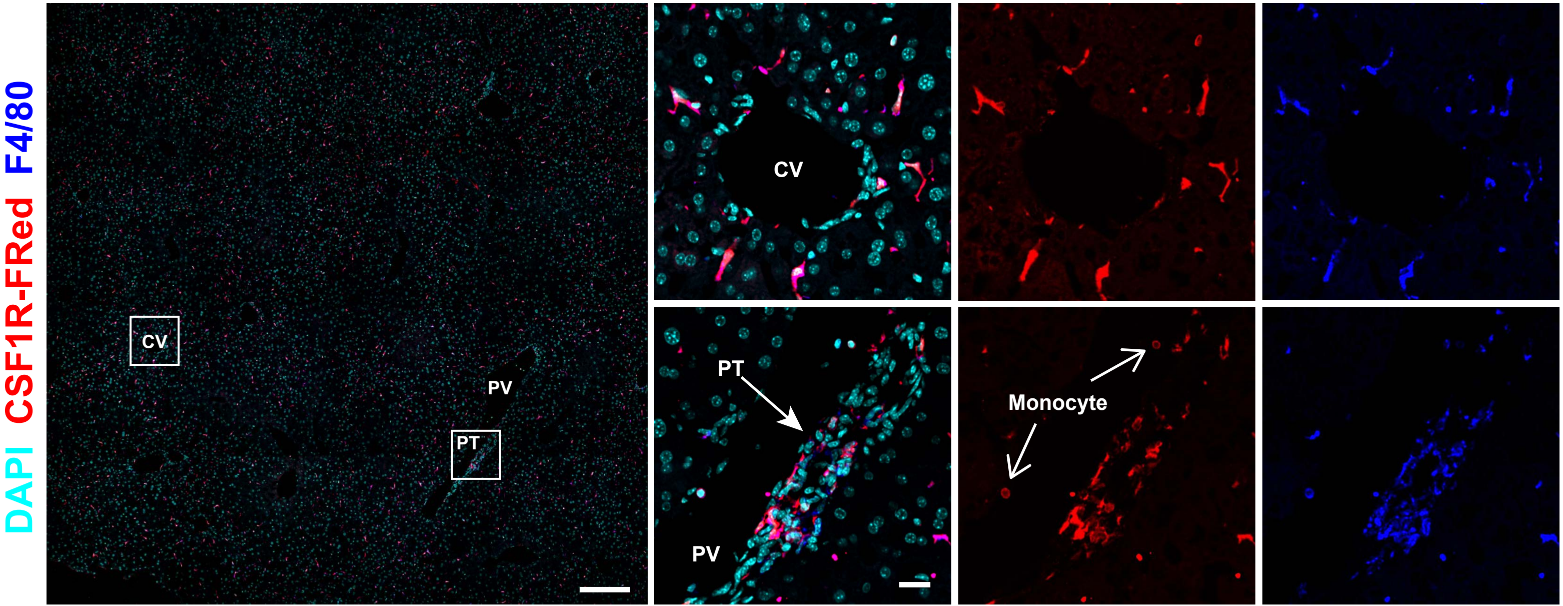


**Figure S4**

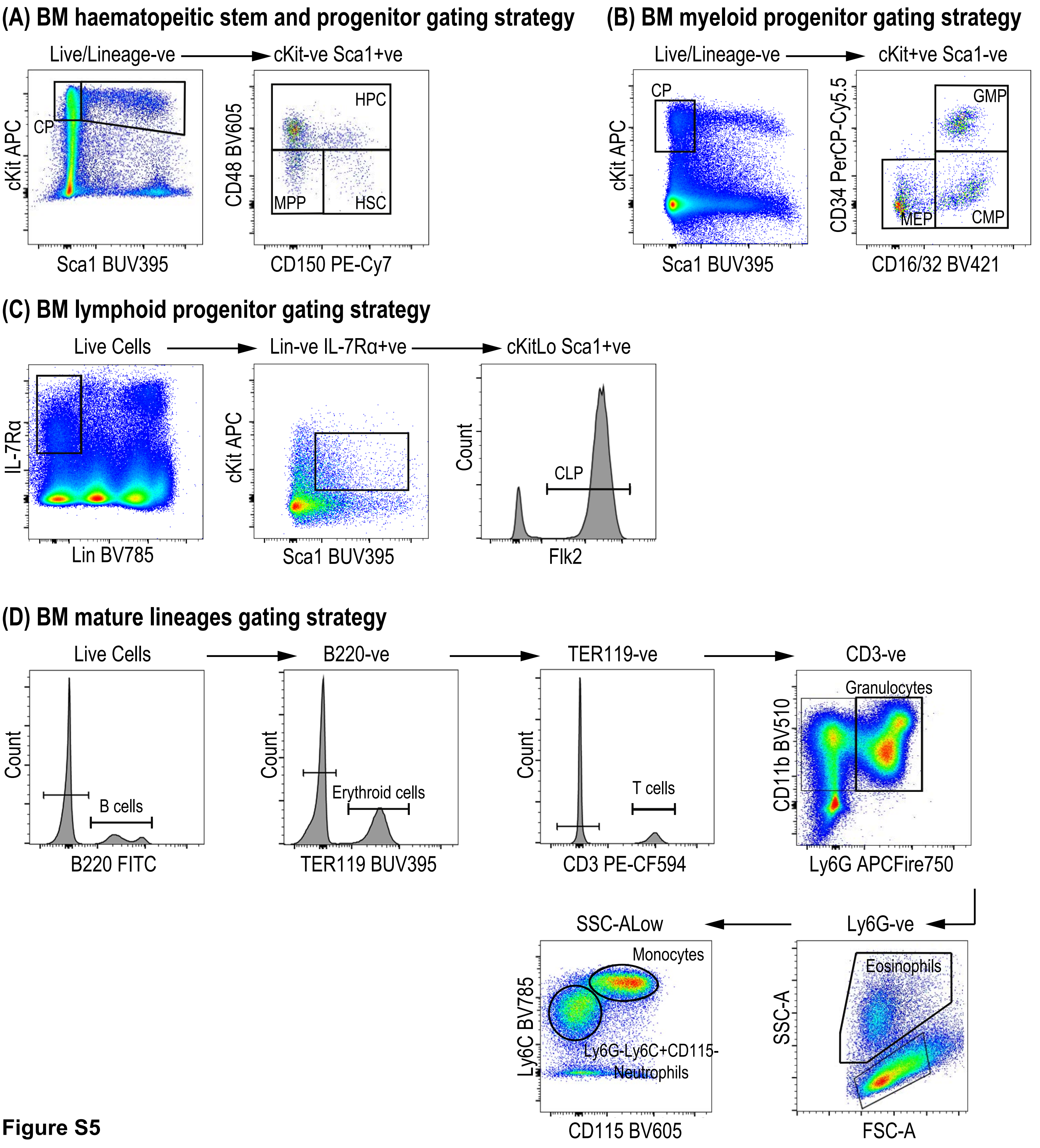
(A) BM haematopeitic stem and progenitor gating strategy
Live/Lineage-ve
cKit-ve Sca1+ve
CP
cKit APC
Sca1 BUV395
HPC
MPP
HSC
CD48 BV605
CD150 PE-Cy7
(B) BM myeloid progenitor gating strategy
Live/Lineage-ve
cKit+ve Sca1-ve
CP
cKit APC
Sca1 BUV395
GMP
MEP
CMP
CD34 PerCP-Cy5.5
CD16/32 BV421
(C) BM lymphoid progenitor gating strategy
Live Cells
Lin-ve IL-7Rα+ve
cKitLo Sca1+ve
IL-7Rα
Lin BV785
cKit APC
Sca1 BUV395
CLP
Count
Flk2
(D) BM mature lineages gating strategy
Live Cells
B220-ve
TER119-ve
CD3-ve
Count
B cells
B220 FITC
Erythroid cells
TER119 BUV395
T cells
CD3 PE-CF594
Granulocytes
CD11b BV510
Ly6G APCFire750
SSC-ALow
Ly6G-ve
Monocytes
Ly6G-Ly6C+CD115-
Neutrophils
Ly6C BV785
CD115 BV605
Eosinophils
SSC-A
FSC-A

**Figure S5**

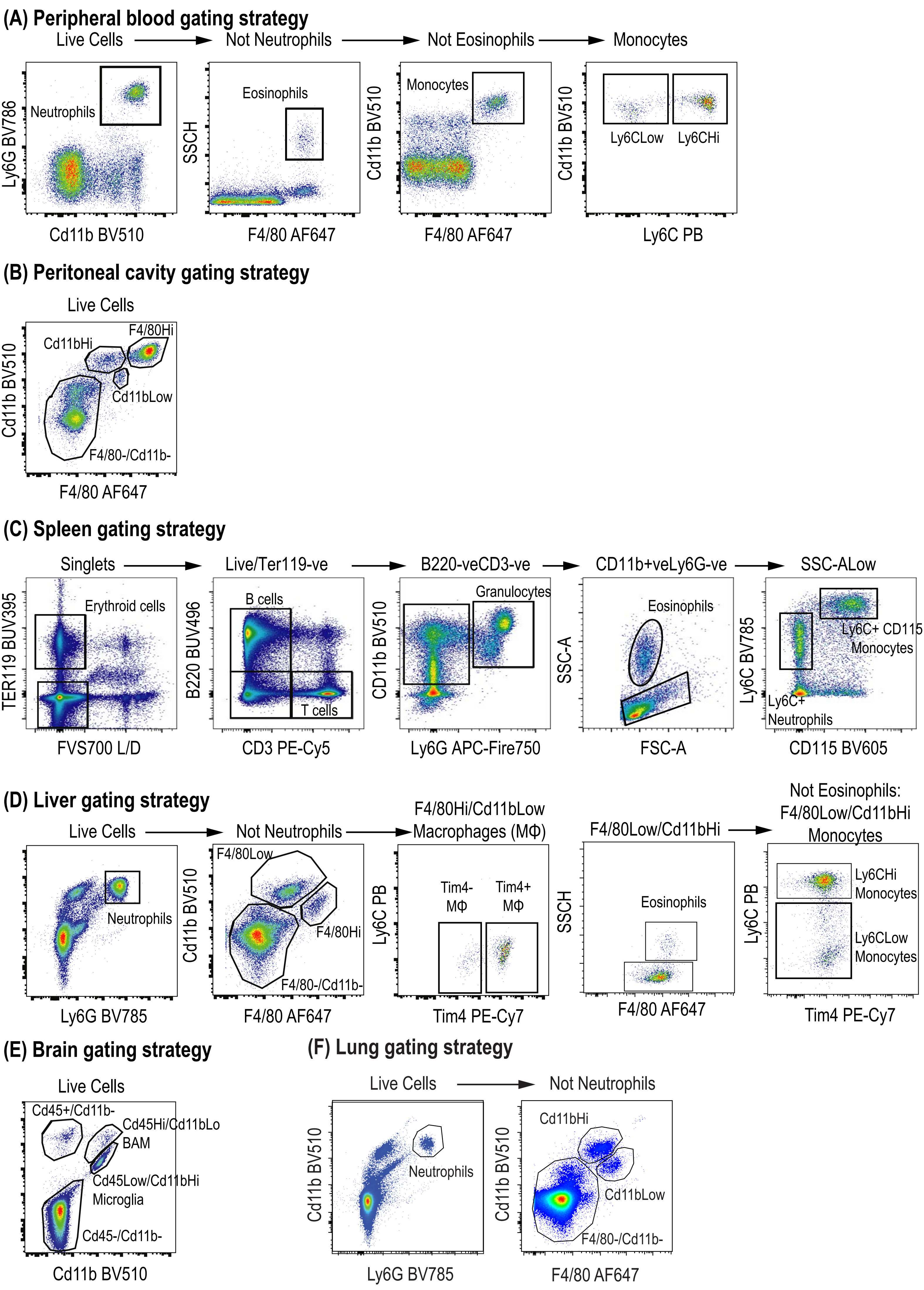
(A) Peripheral blood gating strategy
Live Cells
Not Neutrophils
Not Eosinophils
Monocytes
Neutrophils
Ly6G BV786
Cd11b BV510
Eosinophils
SSCH
F4/80 AF647
Monocytes
Cd11b BV510
F4/80 AF647
Ly6CLow
Ly6CHi
Cd11b BV510
Ly6C PB
(B) Peritoneal cavity gating strategy
Live Cells
Cd11bHi
F4/80Hi
Cd11bLow
F4/80-/Cd11b-
Cd11b BV510
F4/80 AF647
(C) Spleen gating strategy
Singlets
Live/Ter119-ve
B220-veCD3-ve
CD11b+veLy6G-ve
SSC-ALow
Erythroid cells
TER119 BUV395
FVS700 L/D
B cells
T cells
B220 BUV496
CD3 PE-Cy5
Granulocytes
CD11b BV510
Ly6G APC-Fire750
Eosinophils
SSC-A
FSC-A
Ly6C+ CD115
Monocytes
Ly6C+
Neutrophils
Ly6C BV785
CD115 BV605
(D) Liver gating strategy
Live Cells
Not Neutrophils
F4/80Hi/Cd11bLow
Macrophages (MΦ)
F4/80Low/Cd11bHi
Not Eosinophils:
F4/80Low/Cd11bHi
Monocytes
Neutrophils
Ly6G BV785
F4/80Low
F4/80Hi
F4/80-/Cd11b-
Cd11b BV510
F4/80 AF647
Tim4-
MΦ
Tim4+
MΦ
Ly6C PB
Tim4 PE-Cy7
Eosinophils
SSCH
F4/80 AF647
Ly6CHi
Monocytes
Ly6CLow
Monocytes
Ly6C PB
Tim4 PE-Cy7
(E) Brain gating strategy
Live Cells
Cd45+/Cd11b-
Cd45Hi/Cd11bLo
BAM
Cd45Low/Cd11bHi
Microglia
Cd45-/Cd11b-
Cd11b BV510
(F) Lung gating strategy
Live Cells
Not Neutrophils
Neutrophils
Cd11b BV510
Ly6G BV785
Cd11bHi
Cd11bLow
F4/80-/Cd11b-
Cd11b BV510
F4/80 AF647

**Figure S6**

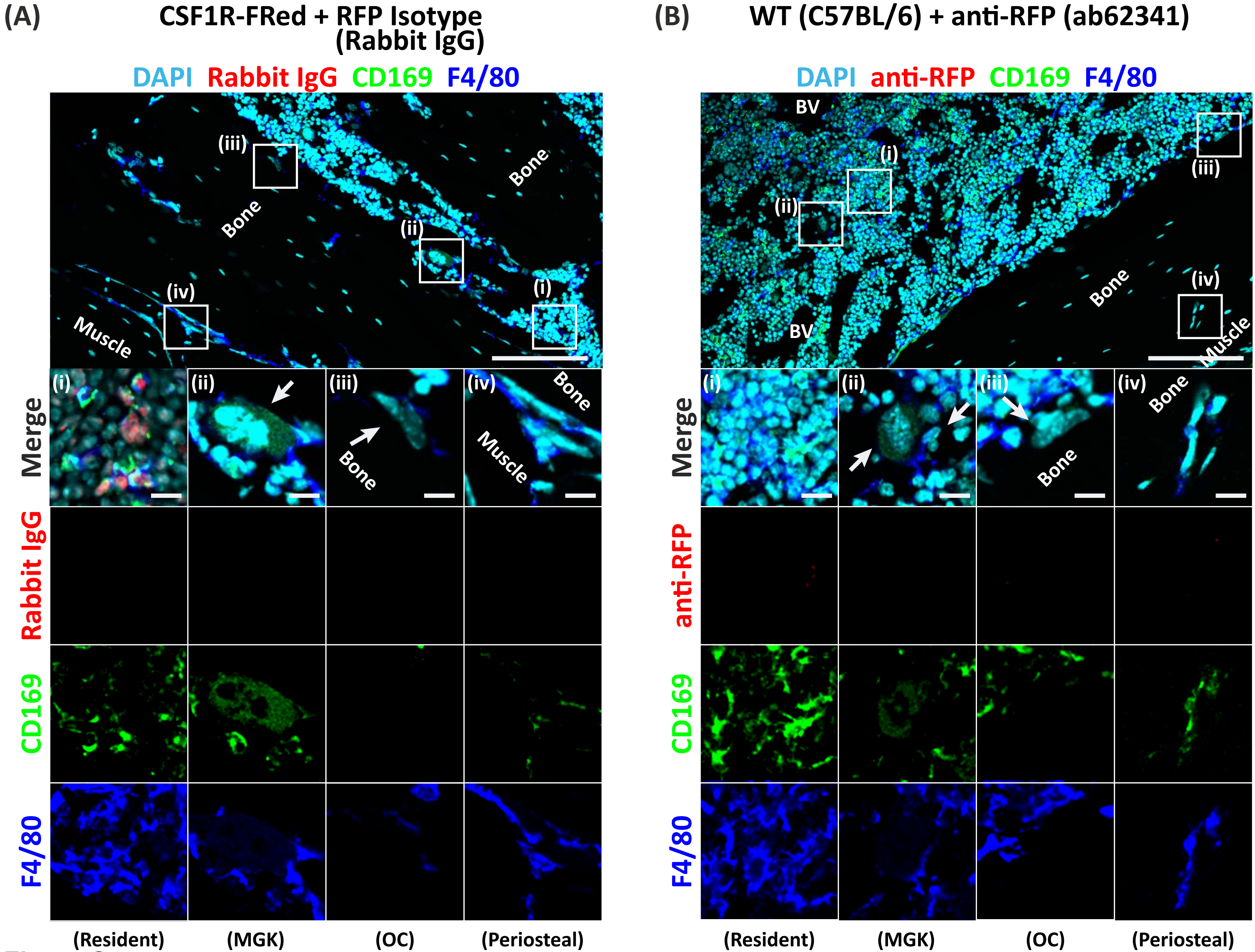
(A)
CSF1R-FRed + RFP Isotype (Rabbit IgG)
DAPI Rabbit IgG CD169 F4/80
Bone
Bone
Muscle
(i)
(ii)
(iii)
(iv)
Merge
Rabbit IgG
CD169
F4/80
(Resident)
(MGK)
(OC)
(Periosteal)
(B)
WT (C57BL/6) + anti-RFP (ab62341)
DAPI anti-RFP CD169 F4/80
BV
BV
Bone
Muscle
Merge
anti-RFP
CD169
F4/80
(Resident)
(MGK)
(OC)
(Periosteal)

Figure S7
